# RT-nested and interfering-Primer PCR reveal prevalent isoform-specific A-to-I RNA editing in neuronal genes

**DOI:** 10.64898/2026.05.15.725286

**Authors:** Ziyi Wang, Yang Ni, Wanzhi Cai, Hu Li, Yuange Duan

**Author notes:** Corresponding author: Yuange Duan, Ph. D. Corresponding author: Hu Li, Ph. D. Other authors’ emails: Ziyi Wang, Yang Ni, Wanzhi Cai.

## Abstract

**Background:** Metazoan adenosine-to-inosine (A-to-I) mRNA editing temporospatially diversifies the neuronal transcriptome and proteome. The limited read length from next-generation sequencing (NGS) constrains the quantification of the potentially differential editing levels across different splicing isoforms, restricting our understanding of the extent to which RNA editing contributes to molecular diversity and its interplay with splicing.

**Methods:** We employed reverse transcription nested PCR (RT-nPCR) and developed a novel interfering-Primer PCR (iPrimer PCR) technique to distinguish different transcripts of any gene. We selected multiple essential genes exhibiting RNA editing in coding sequences (CDSs) or untranslated regions (UTRs) for isoform-specific amplification and Sanger sequencing.

**Results:** Nine different *Adar* isoforms together with pre-mRNA had distinct editing levels at the S>G auto-recoding site, which was predicted to have isoform-specific effects on catalytic activities. Although pre-mRNA editing might exert isoform-dependent promotion/suppression of splicing, closely located editing sites, such as those in neuronal genes *qvr* and *stj*, still exhibited high correlation in editing levels due to co-editing. iPrimer strategy further discovered differential recoding levels between the long/short 3’UTR isoforms of gene *jef*.

**Conclusions:** We provide the first comprehensive solution for isoform-specific PCR amplification of any gene, enabling quantification of RNA editing level of different isoforms. Our results offer insights into how RNA editing interplays with splicing, and highlight its complicated role in expanding molecular diversity.

**Graphical abstract:** 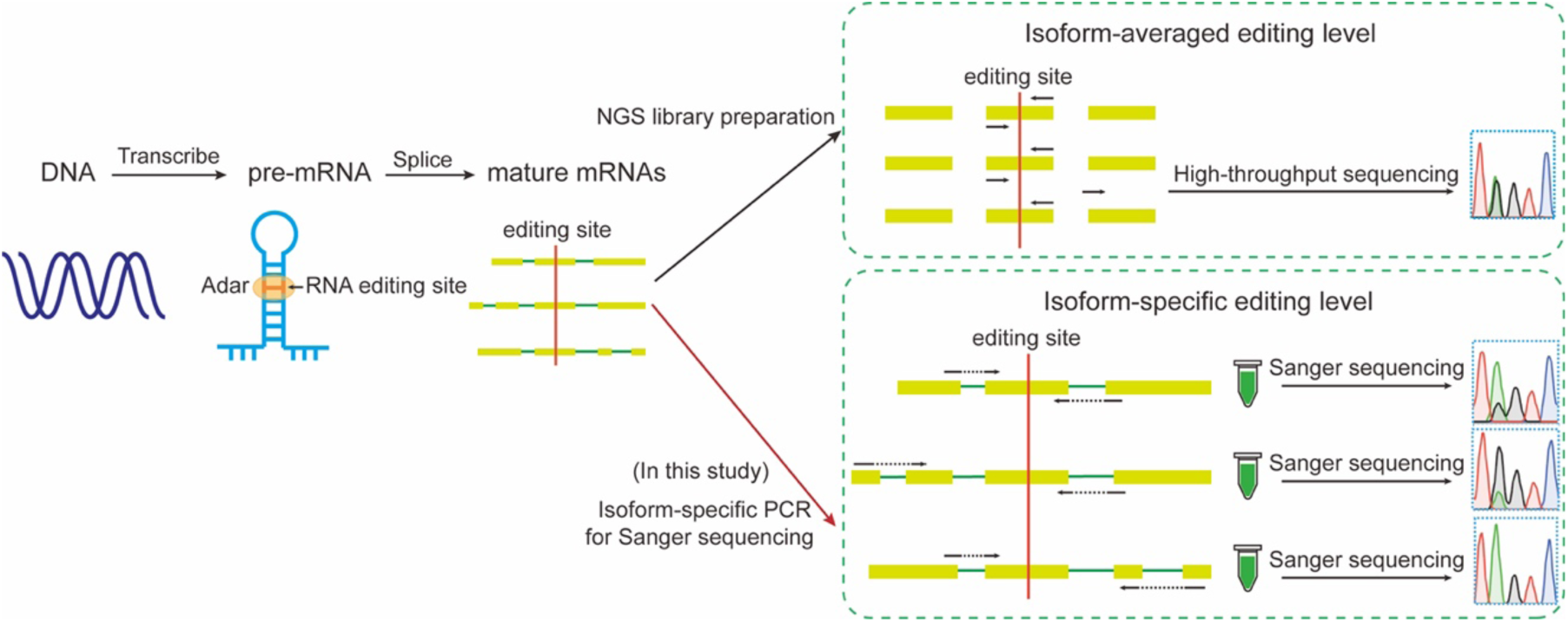

We developed isoform-specific PCR followed by Sanger sequencing, and achieved the quantification of differential RNA editing levels in different transcripts of a gene.

## Introduction

The maturation of eukaryotic RNA involves a series of tightly regulated processing events, including 5’ capping, 3’ polyadenylation, alternative splicing, methylation, and RNA editing [1]. These molecular processes collectively enable precise post-transcriptional regulation without altering the genomic sequence, thereby conferring remarkable plasticity that allows organisms to adapt dynamically to intrinsic physiological demands and complex environmental changes [2, 3].

Adenosine-to-inosine (A-to-I) mRNA editing is a widespread post-transcriptional modification in eukaryotes, catalyzed by the adenosine deaminase acting on RNA (ADAR) family of enzymes [4, 5]. ADARs recognize double-stranded RNA (dsRNA) structures and deaminate specific adenosines to inosines, which are interpreted as guanosines during translation [6–9]. The consequences of RNA editing are diverse, including alterations in protein coding sequences, modulation of RNA secondary structures, and changes in splicing patterns [10–12]. Notably, ADARs exhibit extensive functional interplay with the splicing machinery, as evidenced by observations that knockdown of *ADAR* genes leads to alterations in the splicing patterns of specific transcripts [13].

Alternative splicing generates multiple transcript isoforms from a single gene [14], expanding proteomic diversity and physiological adaptability [15]. However, whether RNA editing efficiency varies across different splicing isoforms of the same gene remains poorly understood. With the advent of next-generation sequencing (NGS), A-to-I RNA editing can be identified and analyzed at the transcriptome-wide level [16, 17]. However, the short read length of NGS technologies limits the ability to resolve distinct isoforms of the same gene, thereby constraining the quantification of isoform-specific editing events. Although Sanger sequencing-based approaches have been used to quantify RNA editing levels [18], their low throughput [19] and reliance on primers spanning specific exon-exon junctions restrict their applicability to genes with relatively few transcript variants [20], while being inadequate for genes with complex isoform repertoires (e.g., more than four transcripts). Other approaches, such as length-resolved PCR, are limited by sequence similarity among isoforms and experimental conditions, making it difficult to distinguish products with similar sizes or sequences using gel electrophoresis [21, 22]. For long-read third-generation sequencing, although it might be a powerful tool in distinguishing different transcript isoforms, its low-throughput nature prevents it from accurate quantification of editing level (which usually requires 100× reads for each single transcript in order to calculate the editing proportion).

Reverse transcription nested PCR (RT-nPCR) combines cDNA synthesis with nested amplification and has been widely used to improve the sensitivity and specificity of low-abundance RNA detection [23]. In this study, we leveraged RT-nPCR by designing multiple primer pairs spanning distinct exon-exon junctions and performing two or more rounds of amplification to obtain separated products for individual transcript isoforms. Furthermore, to address cases in which one transcript is fully encompassed by another (e.g., “UTR isoforms” as defined below), we developed an interfering-Primer PCR (iPrimer PCR) strategy. This method employs specifically designed iPrimers to suppress amplification of longer non-target transcripts, thereby ensuring selective amplification of shorter target isoforms (**Figure 1**). Collectively, this approach enables isoform-specific amplification of any given genes, further allowing quantitative assessment of RNA editing levels at isoform resolution.

**Figure 1.**
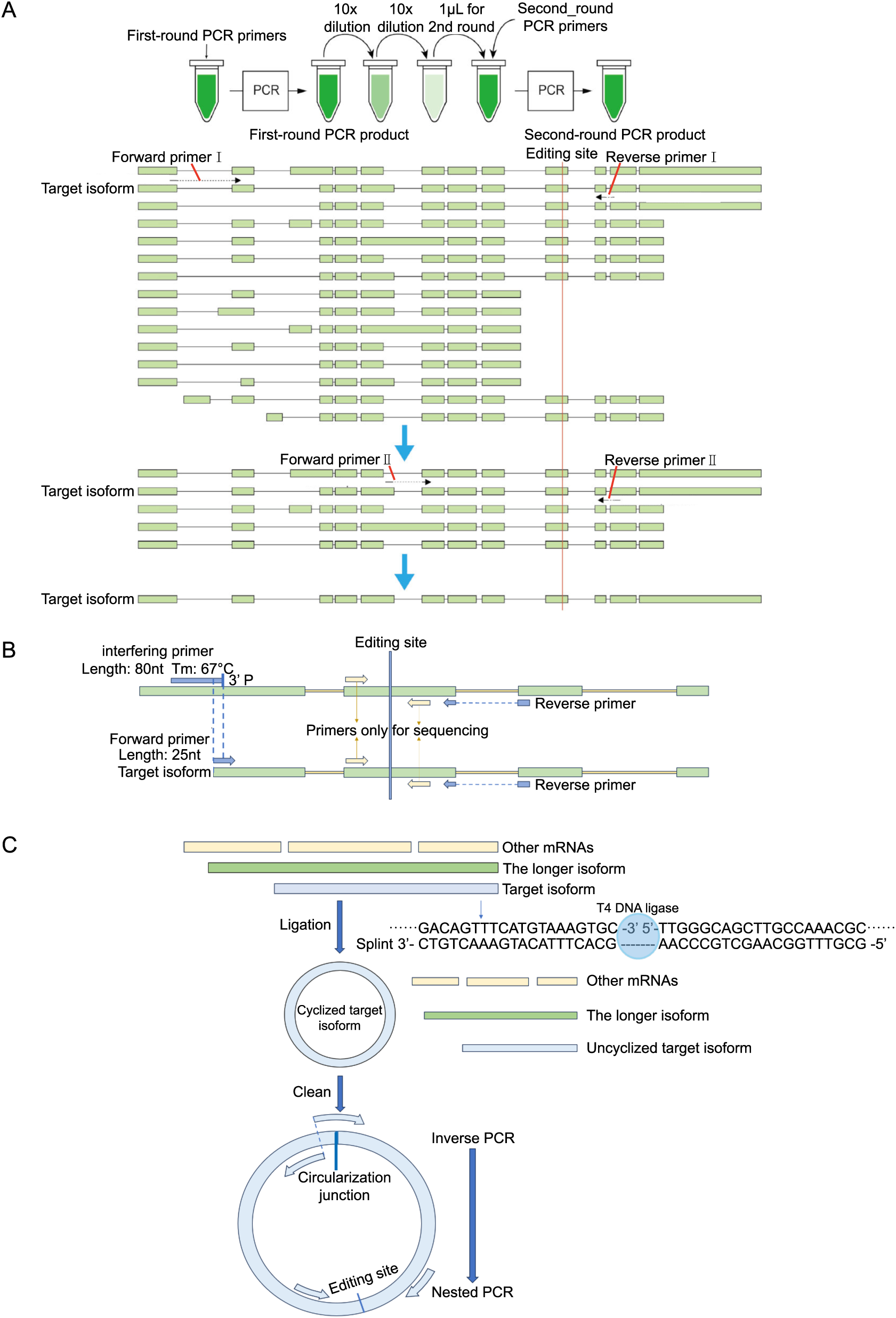
Experimental workflow for isoform-specific PCR amplification used in this study. (A) Take *Adar* gene for instance. We isolate individual *Adar* isoforms using RT-nPCR. The same strategy applies to other transcripts. Multiple primer pairs spanning exon-exon junctions were designed based on transcript sequences, and successive rounds of amplification were performed using different primer sets to obtain high-yield target isoform products. Between each round, PCR products were diluted 100-fold and used as templates for the subsequent amplification. Dilution does not affect the determination of editing levels, but only influences amplification success rate, and can be flexibly adjusted based on enzyme activity, template concentration, and other experimental conditions. For clarity, exon lengths are not shown to scale relative to introns. (B) Schematic illustration of the iPrimer PCR strategy. Interfering primers (iPrimers), designed to suppress amplification of known longer transcript variants, were introduced into the PCR system to ensure selective amplification of target shorter isoforms. (C) Circularized cDNA combined with inverse PCR enables specific amplification of short isoforms within UTR variants. Total cDNA was circularized to generate unique junction sites in short transcripts, which were subsequently targeted for selective amplification by inverse PCR.

Using this strategy, we quantified RNA editing levels across all isoforms for several genes, including *Adar* and *Vps13* (harboring single recoding site), *qvr* and *stj* (containing clustered recoding sites), and *jef* (expressing alternative UTR isoforms). We observed stable differences in recoding levels among isoforms in genes *Adar*, *stj*, and *jef*, suggesting that distinct transcript variants may have differential functional requirements for RNA editing. In contrast, genes such as *Vps13* exhibited variable editing levels across different isoforms but maintained stable editing level for pooled transcripts. Gene *qvr* showed relatively consistent editing across isoforms. These findings indicate that pre-mRNA editing influences the probability of alternative splicing, resulting in differential editing patterns among mature mRNAs. Such isoform-specific regulation may enable RNA editing to fine-tune transcript function and contribute to precise and flexible control of physiological processes. For the first time, we present a comprehensive solution for isoform-specific PCR amplification applicable to all transcripts of a gene, enabling quantitative assessment of RNA editing at isoform resolution. Our findings provide new insights into the interplay between RNA editing and splicing, and underscore its complex role in expanding molecular diversity.

## Results

### Isoform-specific RNA editing at the *Adar* S>G auto-recoding site

The *Drosophila* Adar protein consists of two N-terminal double-stranded RNA-binding domains (dsRBDs) and a C-terminal catalytic deaminase domain (ADEAMc) (**Figure 2A**) [24]. According to NCBI annotations, the *Adar* gene produces 15 different isoforms: eight encode proteins containing both a complete catalytic domain and two intact dsRBDs, five contain truncated catalytic domains but retain two dsRBDs, and two lack the catalytic domain and possess only a single dsRBD.

**Figure 2.**
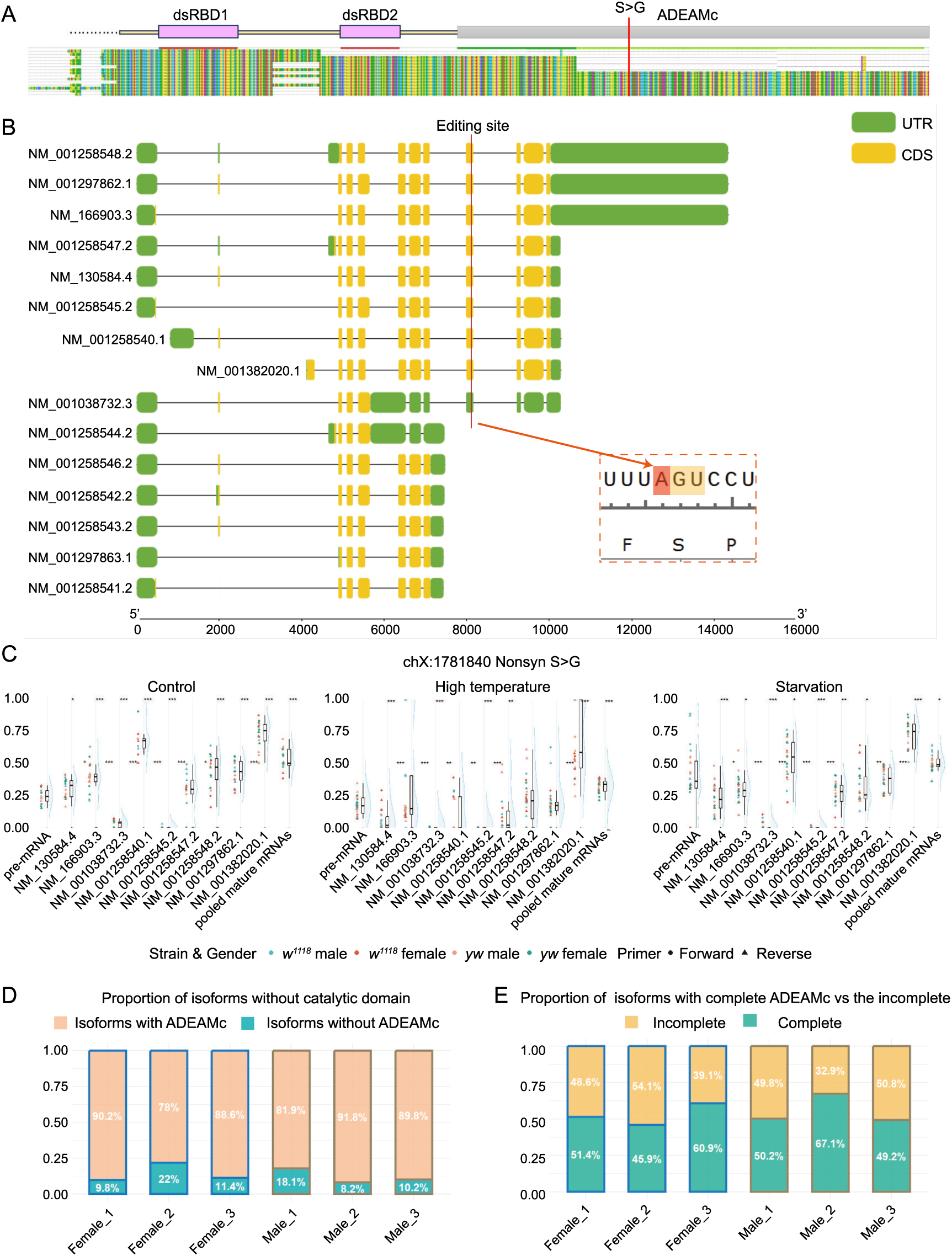
Gene structure, RNA editing levels, and expression profiles of *Adar* isoforms. (A) Domain architecture of the *Drosophila* Adar protein, consisting of two dsRBDs and one catalytic ADEAMc domain. Protein sequences and domain annotations were obtained from NCBI. (B) The *Adar* gene produces 15 different mature isoforms, of which nine contain the S>G editing site. Eight isoforms encode a complete catalytic domain, five encode truncated catalytic domains, and two lack the catalytic domain entirely. (C) RNA editing levels at the *Adar* chrX:1781840 (S>G auto-recoding) site across nine transcript isoforms, pre-mRNA, and pooled mature mRNA. Each isoform includes up to 24 data points (2 strains × 2 sexes × 2 primer sets × 3 biological replicates), with some missing values. Scatter plots show individual data points, with colors indicating strain-sex combinations and shapes indicating primer types. Boxplots summarize distribution statistics (median and interquartile range), and ridge plots display smoothed density distributions (scale = 0.3). Asterisks above indicate significance levels from Wilcoxon rank-sum tests comparing each isoform to pre-mRNA, while asterisks between groups indicate significance between adjacent mature mRNA isoforms: *P* < 0.05 (*), *P* < 0.01 (*), *P* < 0.001 (***). (D) Relative expression proportions of transcript isoforms lacking the catalytic domain, measured by RT-qPCR in male and female *w^1118^* flies (three biological replicates each). Orange indicates isoforms containing a catalytic domain (complete or truncated), and blue indicates isoforms lacking the catalytic domain. (E) Relative expression proportions of isoforms with complete *versus* truncated catalytic domains, measured by RT-qPCR in male and female *w^1118^* flies (three biological replicates each). Yellow indicates isoforms with complete catalytic domains, and green indicates isoforms with truncated catalytic domains.

Notably, the *Drosophila Adar* gene harbors a conserved auto-recoding site (chrX:1781840), where RNA editing converts a serine codon (AGT) into a glycine codon (GGT) [25]. Although only eight isoforms contain a complete catalytic domain, a total of nine transcripts include this editing site, because transcript NM_001038732.3 has this editing event in its 3’UTR (**Figure 2B**). The edited Gly isoform exhibits reduced enzymatic activity, forming a negative feedback regulatory loop [26]. However, whether editing level at this S>G site exhibits isoform specificity remains unclear.

To address this, we analyzed *Adar* auto-recoding levels in male and female *Drosophila melanogaster* of the *w^1118^* and *yw* strains using Sanger sequencing. Editing levels were quantified for *Adar* pre-mRNA, nine mature mRNA isoforms, and pooled mature mRNA samples (see **Materials and Methods**). The results showed that editing levels were relatively consistent across strains and sexes, whereas substantial differences were observed among different isoforms (**Figure 2C**). Environmental perturbations altered both the magnitude and variability of editing across isoforms: under elevated temperature conditions, most mature transcripts exhibited a marked decrease in editing levels, whereas starvation had relatively minor effects (**Figure 2C**). These findings indicate that the S>G auto-recoding site differentially modulates the Adar proteins produced from distinct transcript isoforms and may exert isoform-specific effects on alternative splicing.

To further investigate the prevalence of isoform-specific variation at single editing sites, we examined gene *Vps13* (*Vacuolar protein sorting 13*). *Vps13* encodes a protein located to endosomal membrane that is involved in protein homeostasis. In *Drosophila melanogaster*, Vps13 maintains protein homeostasis in the larval and adult brains where the protein is highly expressed [27]. *Vps13* mutants show shortened lifespan, age-associated neurodegeneration, and accumulation of ubiquitylated proteins [28], but the database-recorded RNA recoding site in this gene has not yet been explored. We noticed that *Vps13* expresses two transcript isoforms that differ in their 5’UTRs (internal difference, not nested), and both isoforms harbor two distantly located editing sites (∼5632 nt apart). We focused on the highly edited nonsynonymous site at chr2R:7579706 (**Supplementary Figure S1A**). In contrast to *Adar*, editing levels did not differ significantly between *Vps13* isoforms (*P* > 0.05), possibly due to the high variability across different samples. However, editing levels in pooled mature mRNA were relatively stable and less dispersed (**Supplementary Figure S1B-C**). These results suggest that for genes whose isoforms are structurally and functionally similar, such as *Vps13*, isoform-specific editing variability may be less informative, whereas pooled editing levels provide a more robust measure of overall RNA editing activity.

### *Adar* isoforms lacking the catalytic domain are lowly expressed

An important question is whether Adar isoforms lacking part or all of the catalytic domain, such as those with truncated ADEAMc domains or the NM_001038732.3 isoform which completely lacks the catalytic domain, perform physiological functions *in vivo*. From an evolutionary perspective, expression patterns with higher functional reward are preferentially retained. In general, highly expressed isoforms tend to fulfill core physiological roles, whereas lowly expressed isoforms are more likely to be evolutionary noises with limited functional contribution [29].

To investigate this, we quantified the relative expression levels of *Adar* isoforms with partially truncated or completely absent catalytic domains in the *Drosophila w^1118^* strain using RT-qPCR. We found that isoforms lacking the ADEAMc domain were generally expressed at very low levels (**Figure 2D**), suggesting that they are unlikely to play major functional roles. In contrast, isoforms with partially truncated catalytic domains exhibited expression levels comparable to those of isoforms with intact catalytic domains, indicating that they may retain important biological functions (**Figure 2E**).

### Isoform-specific effects of the *Adar* S>G recoding site on protein conformation

To further elucidate the functional consequences of recoding on the Adar enzyme, we extracted a conserved RNA secondary structure surrounding the editing site from *Adar* pre-mRNA as a representative RNA substrate for subsequent computational prediction (**Supplementary Figure S2**). This site exhibits stable editing levels and does not have nearby editing sites. For the editing enzyme itself, structural predictions by AlphaFold3 indicate that the Adar S>G recoding site is located within a loop region of the ADEAMc domain, where it modulates deaminase activity by altering local conformational flexibility. Consistent with previous studies suggesting that this recoding event reduces RNA-binding affinity [26], we found that five out of eight Adar isoforms exhibited a decrease in the number of hydrogen bonds with RNA following editing (Gly form), two showed an increase, and one remained unchanged (**Figure 3**). These differences can be attributed to structural features such as dsRNA unwinding capacity, as well as the number and distribution of hydrogen bond interactions. Additionally, the spatial distance between the catalytic domain and the target editing site in the lowest-energy conformation appears to be a critical determinant.

**Figure 3.**
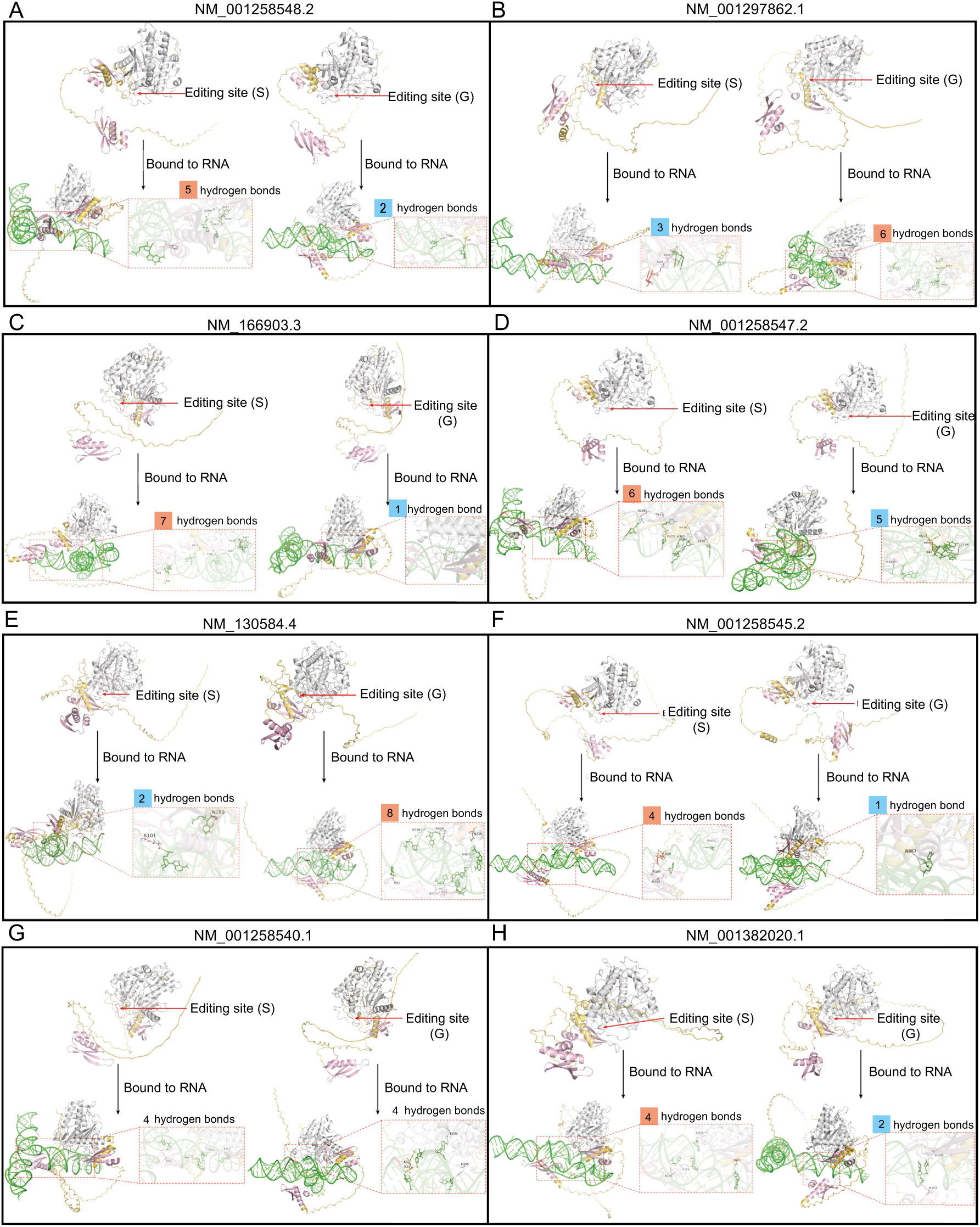
Effects of the *Adar* S>G auto-recoding site on the tertiary structure of the full catalytic Adar protein predicted by AlphaFold3, and analysis of Adar-RNA substrate interactions. (A-H) The S>G site in the Adar protein is located on a loop within the ADEAMc domain, and modulates protein activity by altering the flexible conformation of the catalytic domain. The impact of the S>G editing site varies across different Adar isoforms. These differences are mainly achieved through the formation of distinct flexible structures among transcripts, as well as variations in RNA-binding sites and binding numbers. The number of hydrogen bonds formed between Adar and RNA are indicated. Red numbers denote the conformations with more hydrogen bonds, while blue numbers denote the conformations with fewer hydrogen bonds.

We noticed that the impact of the S>G recoding event varies across isoforms. For example, the protein encoded by NM_001258545.2 displays substantially stronger RNA-binding affinity in the Ser (unedited) form compared to the Gly (edited) form (**Figure 2C** and **Figure 3F**), and correspondingly exhibits extremely low S>G editing levels approaching zero. In contrast, for NM_130584.4, the Gly form increases the number of RNA interaction sites, and this isoform exhibits intermediate editing levels (∼0.2-0.4) (**Figure 2C** and **Figure 3E**). Taken together, these findings demonstrate that the S>G editing event exerts distinct structural and functional effects on different Adar isoforms, underscoring the potential physiological importance of isoform-specific regulation of editing levels.

For Adar isoforms lacking the ADEAMc domain entirely, structural predictions suggest that RNA binding is mediated solely by the remaining dsRBD1 domain. However, these isoforms are unable to unwind double-stranded RNA, and thus cannot perform essential catalytic functions (**Supplementary Figure S3A-B**), consistent with their low expression levels observed by qPCR (**Figure 2D**).

In contrast, isoforms containing partially truncated ADEAMc domains retain the ability to bind RNA and unwind double-stranded structures (**Supplementary Figure S3C-H**), supporting their potential functional relevance, in agreement with qPCR results (**Figure 2E**). Previous studies have proposed that truncated Adar isoforms may perform non-catalytic roles, such as regulation of splicing, and are evolutionarily conserved [30]. Our molecular modeling results provide further support for this hypothesis.

### Editing levels of clustered editing sites are highly correlated across different transcripts

Clustered editing sites refer to regions within RNA molecules where high-density A-to-I editing events occur, typically spanning tens to hundreds of nucleotides [31, 32]. To investigate the editing patterns of clustered editing sites across different transcripts, we selected the *qvr* gene (with two isoforms) and the *stj* gene (with three isoforms) in *Drosophila*, and performed isoform-specific amplification followed by Sanger sequencing to quantify editing levels.

*Drosophila qvr* (*quiver*) encodes a Ly-6 protein that modulates potassium channels and nicotinic acetylcholine receptors, and is extensively recoded at multiple conserved sites [33]. These modifications likely contribute to sleep regulation and ion channel modulation in the nervous system [33–35]. Gene *stj* (*straightjacket*) encodes a voltage-gated calcium channel subunit α2δ3 that stabilizes Cacophony (Cac) α1 subunits at synapses, regulating neurotransmitter release probability [36]. Loss of *stj* causes reduced synaptic transmission, seizure-like activity, and defects in thermal nociception [37, 38]. To date, RNA editing of the *Drosophila stj* has only been recorded in bioinformatic studies without further exploration.

There are eight *qvr* editing sites located within the same hairpin stem, and their editing levels are relatively stable across different sites or different samples (**Supplementary Figure S4**). Notably, four nonsynonymous sites (chr2R:11447601, 11447607, 11447617, and 11447623) maintain relatively high editing levels in mature mRNA (**Supplementary Figure S4**), while three synonymous sites (chr2R:11447606, 11447612, and 11447621) and one nonsynonymous site (chr2R:11447625) exhibit low editing levels approaching zero (**Supplementary Figure S5**). This supports a selective advantage on nonsynonymous editing as multiple previous studies indicated [39, 40].

Six *stj* recoding sites reside in different regions of the dsRNA structure (**Figure 4A** and **4B**). Editing levels at the same site vary substantially across different transcripts. Under elevated temperatures, editing levels fluctuate and generally show varying degrees of reduction across mature mRNAs while the overall editing levels remain narrowly distributed (**Figure 4C-F**; **Supplementary Figure S6A**). There is one site (chr2R:13817124) with editing levels approaching zero (**Supplementary Figure S6B**).

**Figure 4.**
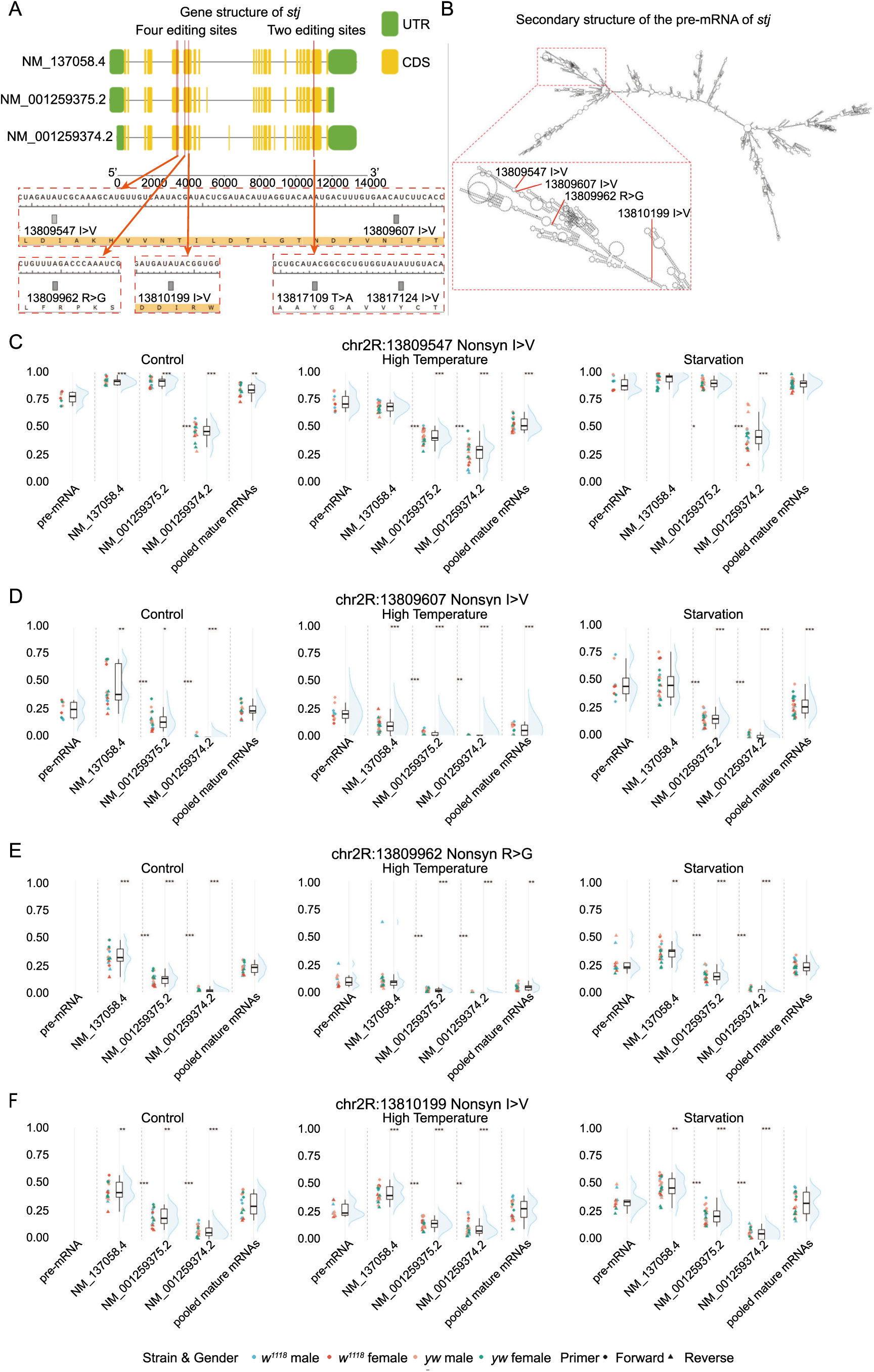
Isoform-specific quantification of *stj* RNA editing levels using Sanger sequencing. (A-B) The first four adjacent editing sites in *stj* are located within different dsRNA structures. (C-G) Editing levels at the same sites vary significantly across different *stj* isoforms (*P* < 0.05), although their distributions remain relatively clustered. The y-axis represents RNA editing levels. Under different experimental conditions, particularly at elevated temperatures, editing levels fluctuate across transcripts. The plotting scheme is consistent with that used for the *Adar* gene.

In clustered editing regions, the spatial proximity of editing sites facilitates sequential editing by Adar proteins. After completing one editing event, Adar is more likely to edit adjacent sites, resulting in correlated editing levels among nearby positions [41]. To examine whether clustered editing sites exhibit similar co-editing patterns across different transcripts of the same gene, we investigated the eight adjacent *qvr* sites and four adjacent *stj* sites across different samples (see **Materials and Methods**). Four clustered *qvr* sites (chr2R:11447601, 11447607, 11447617, and 11447623) show strong correlation in editing levels, suggesting they are frequently edited simultaneously by Adar. Although another four *qvr* sites (chr2R:11447606, 11447612, 11447621, and 11447625) also appear highly coordinated, their editing levels are relatively low and thus might lack clear biological significance (**Figure 5A** and **5B**). For *stj* gene, correlations among sites are notably weaker compared to *qvr* (**Figure 5C-E**), which is consistent with the greater distances between adjacent sites in *stj* relative to those in *qvr*.

**Figure 5.**
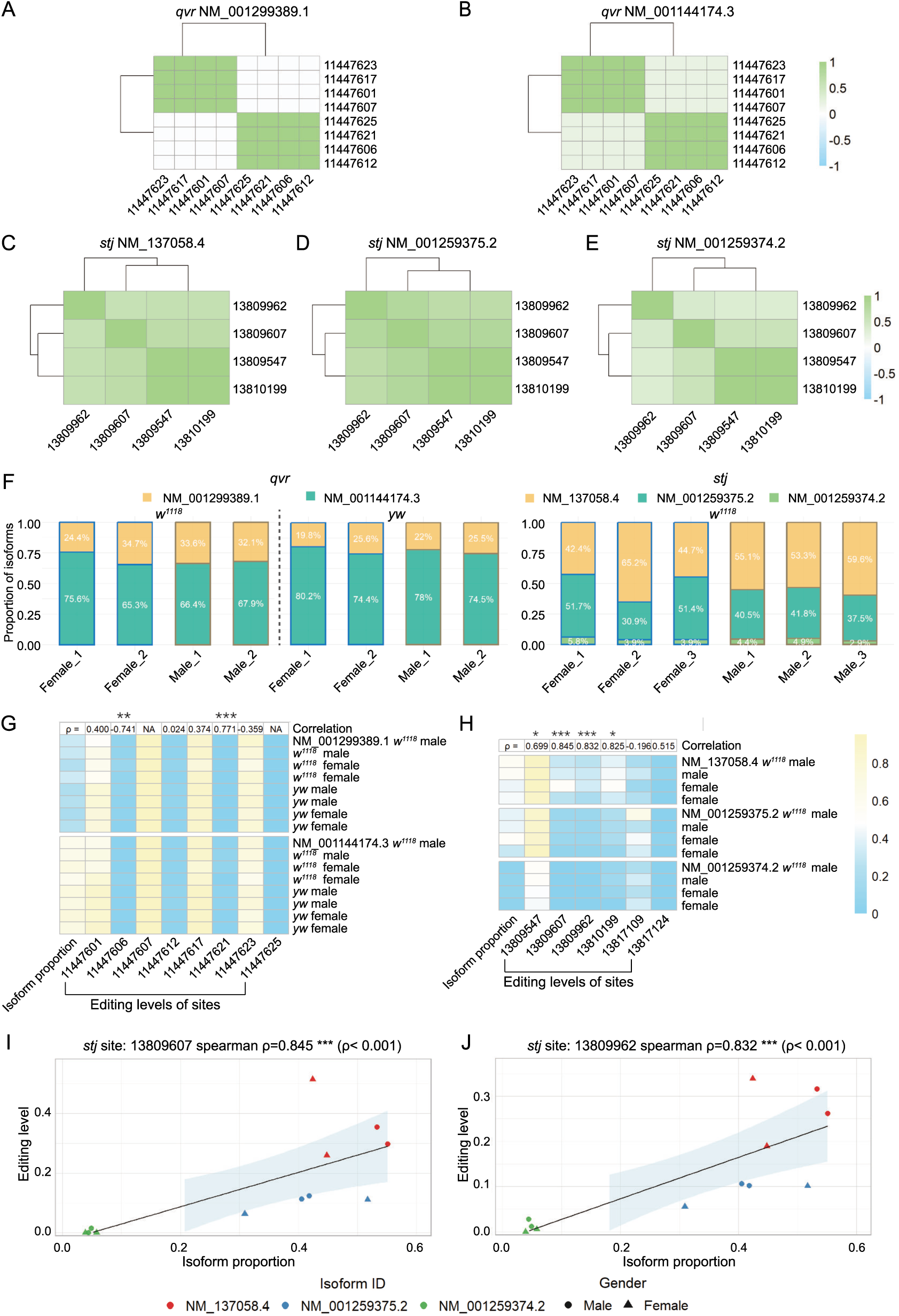
Co-editing patterns of clustered editing sites of qvr and stj, and specific sites show correlations with transcript expression levels. (A-E) Pearson correlation analysis reveals co-editing relationships among editing sites in *qvr* and *stj*. Heatmaps display strong positive correlations between certain site pairs (highlighted in dark green), whereas others appear weakly correlated or independent (shown in light colors). In *qvr*, four sites (chr2R:11447601, 11447607, 11447617, and 11447623) exhibit strong coordination, indicating simultaneous editing. Another four sites (chr2R:11447606, 11447612, 11447621, and 11447625) also appear highly coordinated, their editing levels are low. In *stj*, correlations among sites are weaker than those observed in *qvr*, consistent with the greater distances between adjacent editing sites. (F) Relative expression levels of individual transcripts of *qvr* and *stj* were quantified using RT-qPCR. For *qvr*, two biological replicates were measured for both males and females in the *w^1118^* and *yw* strains. For *stj*, three biological replicates were measured for males and females in the *w^1118^*strain. (due to constraints of 96-well plate layout and limited availability of *yw* samples). (G-H) Heatmaps showing correlations between transcript expression proportions (first column) and editing levels at multiple RNA editing sites (the remaining columns) for *qvr* (left) and *stj* (right). The top panel indicates Spearman correlation coefficients (ρ) between editing levels and transcript expression. Statistical significance is annotated above the coefficients: *P* < 0.05 (*), *P* < 0.01 (**), *P* < 0.001 (***). (I-J) Scatter plots for two *stj* editing sites exhibiting strong and significant correlations (*P* < 0.001) with transcript expression and with sufficiently high editing levels, namely sites chr2R:13809607 (left) and chr2R:13809962 (right). Points are colored by transcript and shaped by sex. Linear regression lines with 95% confidence intervals (light blue shading) are overlaid.

### Correlation between editing level and expression level of transcripts

In the light of evolution, if particular RNA recoding events were beneficial, such events should be selectively favored in highly expressed genes/transcripts as the benefit would be amplified. To preliminarily distinguish which editing sites might be essential, we investigated the correlation between editing level and expression level of transcripts. Using qPCR, we quantified the proportion of each transcript relative to the total expression of the *qvr* (two isoforms) and *stj* (three isoforms) genes (**Figure 5F**). Spearman correlation coefficients were calculated across different samples to assess the relationship between transcript expression levels and RNA editing levels at individual sites.

In the *qvr* gene, there is little to no significant correlation at most sites. For sites chr2R:11447606 and chr2R:11447621, although the correlation is significant, the extremely low editing levels render the correlation uninformative (**Figure 5G**). In contrast, in the *stj* gene, editing levels at recoding sites chr2R:13809547 and chr2R:13810199 show significant positive correlations with transcript expression (*P* < 0.05, **Figure 5H**), whereas editing levels at recoding sites chr2R:13809962 and chr2R:13809607 exhibit even stronger correlations with transcript expression (*P* < 0.001, **Figure 5H-J**). Notably, the above four recoding sites that display significant correlations (*P* < 0.05) correspond exactly to the four adjacent clustered editing sites in *stj*.

### Temperature exerts isoform-specific effects on RNA editing level

Temperature can reshape neuronal and locomotor system functions in animals by modulating RNA editing, thereby enabling adaptation to changing environmental conditions [42, 43]. Previous studies have shown that elevated temperature generally leads to a global reduction in RNA editing levels [44]. Here, we calculated the average editing levels across samples and compared the differences between high-temperature and control groups. Four *qvr* sites (chr2R:11447606, 11447612, 11447621, and 11447625) were excluded from further analysis due to their intrinsically low editing levels.

The results show that, for *Adar*, *qvr*, and *stj*, most transcripts exhibit decreased editing levels under high-temperature conditions, with only a few sites showing slight increases (**Figure 6A-C**). Notably, for the same editing site, the magnitude of this decrease varies substantially across different transcripts, highlighting isoform-specific regulation of RNA editing in response to temperature changes. This also suggests that differences in editing levels among isoforms are dynamically regulatable, allowing organisms to fine-tune splice variants with distinct physiological functions under changing environmental conditions.

**Figure 6.**
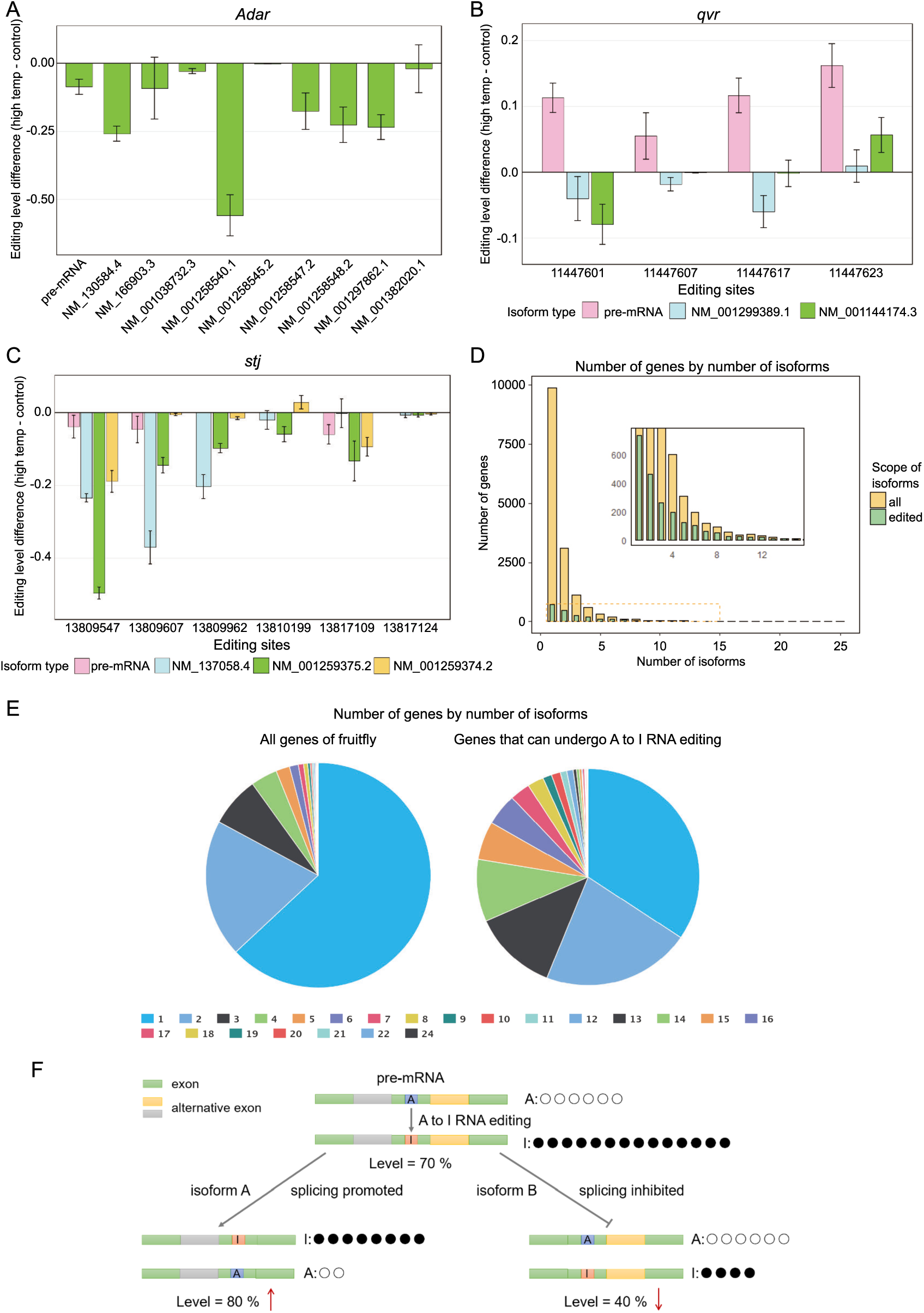
Temperature-sensitive RNA editing events and the variation in editing levels among different mRNA isoforms. (A-C) Differences in mean editing levels between high-temperature and control groups vary significantly across different transcripts. Bars above the x-axis indicate increased editing levels under high temperature, whereas bars below the x-axis indicate decreased editing levels. Error bars represent the standard error of the mean (SEM) of the differences. (D) Genes undergoing RNA editing tend to have a greater number of transcript isoforms. The yellow bars represent the distribution of all *Drosophila* genes according to transcript number, while the nested green bars represent genes with RNA editing events. Genes with multiple transcripts tend to possess RNA editing. (E) Pie charts showing the following proportions. Among all *Drosophila* genes, more than half produce only a single transcript. In contrast, among genes with A-to-I RNA editing events, more than half generate multiple transcripts. Pie chart was generated using the OmicShare platform. (F) A schematic diagram showing the interplay between pre-mRNA editing and alternative splicing. When RNA editing promotes the formation of a specific splicing isoform, the editing level in that transcript is higher than in pre-mRNA; conversely, when RNA editing inhibits the formation of a splicing isoform, its editing level is lower than that in pre-mRNA. Circles represent the number of transcripts of each type in the illustrated examples.

Importantly, while mature mRNA generally shows reduced editing levels at elevated temperatures, pre-mRNA exhibits much smaller changes, and in the case of *qvr*, even a general increase in pre-mRNA editing levels is observed (**Figure 6B**). Previous studies have proposed that increased temperature suppresses A-to-I RNA editing globally, mainly due to reduced stability of RNA secondary structures and regulation of *Adar* expression [45]. However, the relatively stable (or even increased) editing levels observed in pre-mRNA in our study cannot be readily explained by existing models. This phenomenon may involve complex interactions among temperature, RNA editing, and alternative splicing which warrant further investigation.

### RNA editing may differentially promote or inhibit isoform-specific splicing

Both RNA editing and alternative splicing can enhance molecular diversity at transcript level, yet whether these two mechanisms are mutually enriched or mutually excluded remains unclear at genome-wide level. To preliminarily answer this question, we quantified the number of isoforms for genes undergoing RNA editing and compared this distribution with the baseline of all genes in the *Drosophila melanogaster* genome. We found that genes with RNA editing events tend to have a greater number of isoforms, and the proportion of edited genes increases with isoform number (**Figure 6D-E**). This observation suggests a potential positive association between RNA editing and alternative splicing.

Previous studies have shown that RNA editing can regulate alternative splicing through multiple mechanisms, including direct alteration of splice site sequences, modification of splicing regulatory elements, and coordinated editing near intronic splice sites [11, 46, 47]. Using splice site prediction tools from the Berkeley *Drosophila* Genome Project, we analyzed splice donor and acceptor sites in *Adar*, *Vps13*, *qvr*, and *stj*. None of the editing sites examined were located within canonical splice motifs, and sequence alterations at these positions did not affect predicted splicing patterns. This indicates that RNA editing in these genes does not regulate splicing by directly modifying splice site sequences.

It was reported that more than 95% of A-to-I RNA editing events occur in chromatin-associated RNA prior to polyadenylation [11], suggesting that RNA editing predominantly takes place before the completion of splicing and formation of mature mRNA. Given that the editing sites analyzed in *Adar* and *stj* exhibit stable and significant differences in editing levels across transcripts, this implies that RNA editing events at the pre-mRNA level differentially influence the alternative splicing process of distinct isoforms. Moreover, previous studies have demonstrated discrepancies between editing levels in pre-mRNA and mature mRNA, attributed partly to the influence of RNA editing on splicing kinetics or the interference with splicing by ADAR binding [48]. Accordingly, when RNA editing promotes the formation of a specific transcript, the editing level in that transcript is expected to be higher than that in pre-mRNA; conversely, when RNA editing inhibits transcript formation, its editing level is expected to be lower than that in pre-mRNA (**Figure 6F**).

To quantify this effect, we calculated the differences between editing levels in mature mRNA transcripts and those in pre-mRNA from the same samples, using these differences as indicators of the promoting or inhibitory effects of RNA editing on the tendency of isoform formation (**Figure 7A-C**).

**Figure 7.**
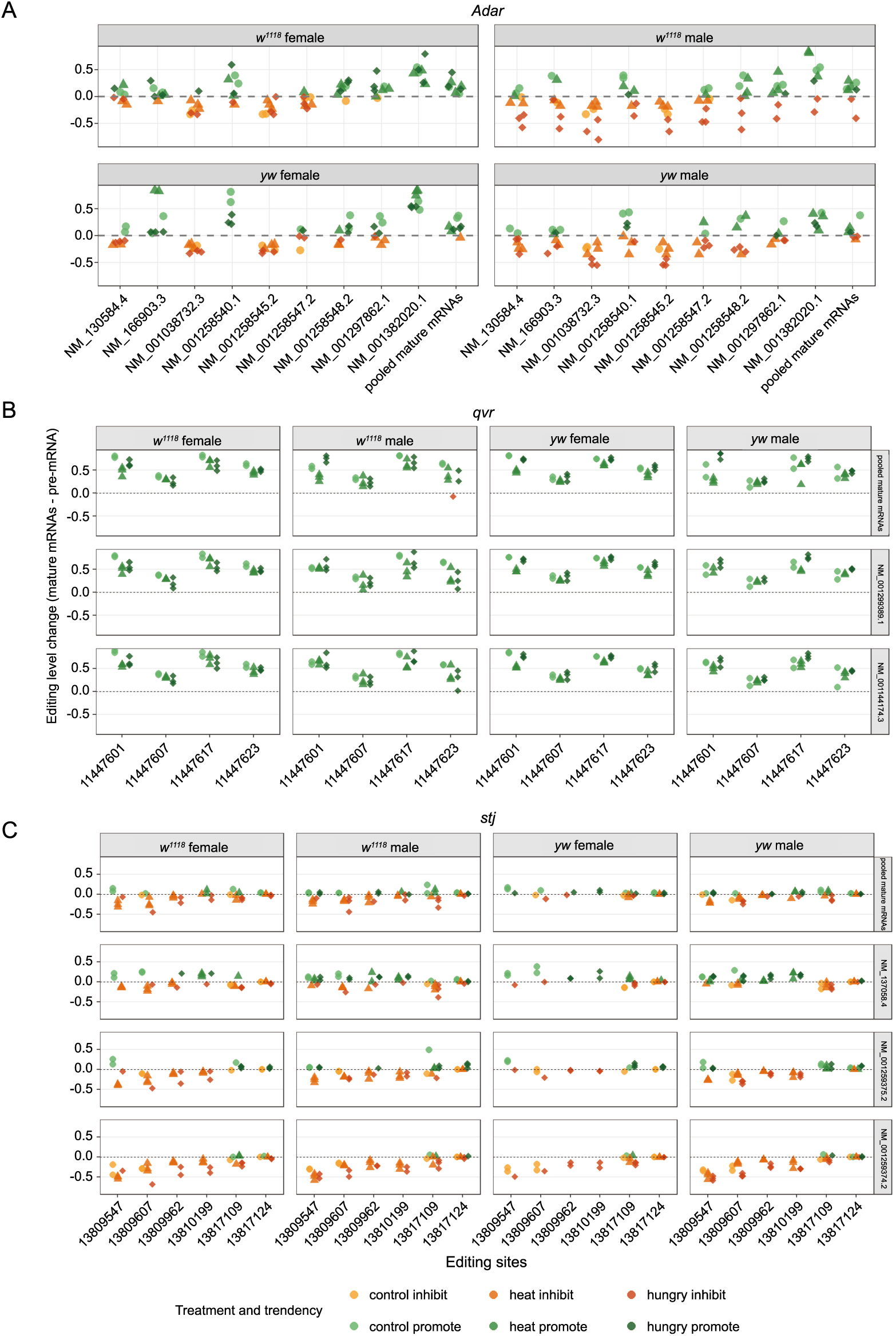
Differences between editing levels in mature mRNA transcripts and pre-mRNA reflect the promoting or inhibitory effects of RNA editing on transcript-specific splicing. (A) Differences between S>G auto-recoding levels of each *Adar* isoform and pre-mRNA. The x-axis represents different transcripts, and the y-axis represents the magnitude of the difference. Each panel corresponds to flies of the same strain and sex. Yellow dots indicate inhibitory effects of RNA editing on splicing, whereas green dots indicate promoting effects. Different point shapes represent different experimental conditions. (B-C) Differences between editing levels of different isoforms and pre-mRNA for *qvr* and *stj*. The x-axis represents genomic coordinates of editing sites. Each panel represents different transcripts from flies of the same strain and sex. Other visualization conventions are consistent with those used for the *Adar* gene.

In the *Adar* gene, substantial differences in editing level changes are observed across transcripts, indicating that RNA editing influences their splicing processes in distinct ways (**Figure 7A**). Moreover, the direction of these changes (promoting or inhibitory) varies across different culture conditions, strains, and sexes, suggesting that the regulatory effects of RNA editing on splicing are not constant but are modulated by environmental and physiological factors. For the *qvr* gene, editing levels in all transcripts are generally higher than those in pre-mRNA, indicating an overall promoting effect of RNA editing on splicing. However, the magnitude of increase varies substantially across sites, suggesting that different editing sites contribute unequally to splicing regulation (**Figure 7B**).

In the *stj* gene, RNA editing exerts distinct effects on the splicing of its three transcripts. For example, transcripts NM_001259374.2 and NM_001259375.2 predominantly show negative regulation, indicating that pre-mRNA editing tends to inhibit their formation. In contrast, transcript NM_137058.4 exhibits both positive and negative changes. Notably, despite differential performances for distinct isoforms, four adjacent sites (chr2R:13809547, 13810199, 13809962, and 13809607) tend to show similar differences, suggesting that editing at nearby sites may act coordinately to influence transcript processing (**Figure 7C**).

### Amplification of “nested” transcripts using interfering-primer PCR and circularized cDNA approaches

UTR isoforms represent an important form of post-transcriptional regulation. These isoforms arise when a single gene produces transcripts (isoforms) with identical CDS but differing lengths of 5’ or 3’ UTRs, where one isoform is completely embedded in another isoform [49]. The presence of UTR isoforms confers high plasticity to gene expression, primarily by regulating mRNA stability, nuclear export efficiency, and interactions with microRNAs or RNA-binding proteins, thereby modulating gene expression patterns across tissues and developmental stages [50–52].

Gene *jef* (*jet fuel*) in *Drosophila* encodes a protein predicted to enable transmembrane transporter activity, functioning as a major facilitator superfamily (MFS) solute carrier involved in membrane transport [53]. This gene is expressed in adult head, embryonic nervous system, and larval ganglia [54] without the mention of the function of its RNA editing sites.

The *jef* gene expresses two 3’UTR isoforms, both of which contain two adjacent RNA editing sites within the CDS: a nonsynonymous S>G editing site (chr2R:1472210) and a synonymous editing site (chr2R:1472223) (**Figure 8A**). However, conventional methods for measuring editing levels, including the nested RT-PCR approach described above, are insufficient for specifically amplifying the shorter transcript when its sequence is fully embedded within that of a longer transcript.

**Figure 8.**
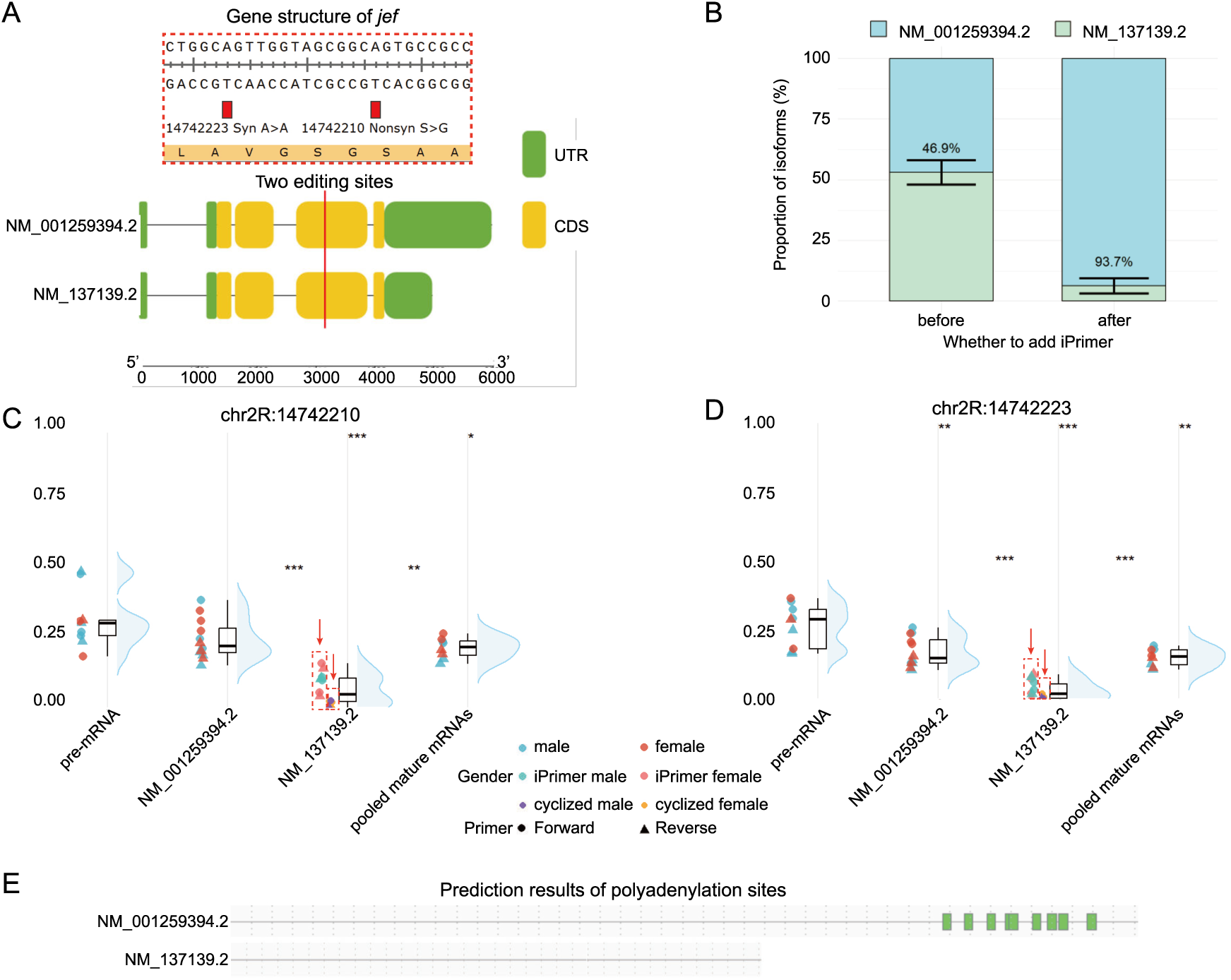
Specific amplification of short UTR isoforms using interfering-primer PCR (iPrimer PCR) and circularized cDNA inverse PCR. (A) Schematic representation of the two UTR isoforms of the *jef* gene and the positions of the analyzed editing sites. The plots were generated using Tbtools-II. (B) Changes in measured expression ratios by qPCR before (left) and after (right) inclusion of the interfering primer in the amplification system targeting the short transcript. The average ratio across samples is indicated by the blue-green dividing line. Error bars represent the standard deviation (SD), with midpoints corresponding to the dividing line. (C-D) Editing levels of *jef* transcripts in the *w^1118^* strain were quantified. Editing levels of NM_137139.2 were measured using iPrimer PCR (blue dots for males, orange for females) and circularized cDNA (purple for males, yellow for females). Visualization follows the same scheme as for the *Adar* gene. (E) Gene model of two *jef* transcripts. NM_137139.2 lacks polyadenylation sites, whereas the additional 3’UTR region in NM_001259394.2 contains multiple polyadenylation sites. Green indicates polyadenylation sites, and gray lines represent other regions. Prediction and visualization were performed using RegRNA 3.0.

To address this challenge, we developed two complementary strategies for double confirmation: (1) interfering-primer PCR (iPrimer PCR) to suppress amplification of the longer transcript, and (2) circularized cDNA followed by inverse PCR to introduce unique junction sites enabling specific amplification of the shorter transcript (see **Materials and Methods**). Furthermore, we combined iPrimer PCR with qPCR to quantify the relative expression levels of the two *jef* UTR isoforms. The inclusion of the interfering primer effectively excluded the non-target long transcript (NM_001259394.2) from the amplification system targeting the short transcript (NM_137139.2), resulting in expression estimates that more accurately reflect the true abundance of NM_137139.2. These results demonstrate the effectiveness of the iPrimer PCR approach. Our results show that the long UTR isoform NM_001259394.2 is highly abundant, whereas NM_137139.2 is expressed at relatively low levels (**Figure 8B**).

Sanger sequencing reveals that both the nonsynonymous editing site (chr2R:1472210) and the synonymous site (chr2R:1472223) exhibit transcript-specific differences in editing levels, with consistently lower editing levels observed in the shorter transcript NM_137139.2 (**Figure 8C** and **8D**). These findings not only support the positive correlation between expression and editing level (an indicator of beneficial RNA editing), but also demonstrate that transcript-specific variation in RNA editing levels can also occur at synonymous sites and in UTR isoforms. Notably, in most cases, the editing levels measured by iPrimer PCR and circularized cDNA methods are highly consistent, which justifies that our novel methodologies are technically sound and robust, while only for isoform NM_137139.2, the circularized cDNA approach yields slightly lower editing levels.

Using RegRNA 3.0, we predicted RNA motifs and functional elements in the two transcripts. The shorter isoform NM_137139.2 lacks polyadenylation sites, whereas the longer 3’UTR in NM_001259394.2 contains multiple polyadenylation sites (**Figure 8E**). This suggests that NM_137139.2 may be retained in the nucleus and potentially degraded as aberrant mRNA, resulting in its low expression level. Again, such correlation of RNA editing level and transcript expression level between these two isoforms aligns with our and others’ notion that potentially beneficial RNA editing might preferentially occur in more essential and highly expressed transcripts.

## Discussion

As an essential epitranscriptomic regulatory layer, A-to-I RNA editing needs to be precisely finetuned across different loci. However, the same genomic locus can be assigned to different locations at distinct splicing isoforms, and thus raising the need to distinguish the editing levels at different isoforms. Differences in A-to-I RNA editing levels among isoforms reflect that RNA editing events occurring at the pre-mRNA level might differentially influence the splicing process or regulation of distinct transcripts. In this study, we employed reverse transcription nested PCR (RT-nPCR) to achieve isoform-specific amplification of multi-transcript genes, followed by Sanger sequencing to quantify isoform-specific editing levels. Furthermore, we developed the interfering-primer PCR (iPrimer PCR) method to enable specific amplification and editing quantification of “nested” transcripts of the same gene. These approaches establish a framework for isoform-specific RNA editing analysis applicable to all genes.

At the S>G auto-recoding site in the *Adar* gene, we observed stable and significant differences in editing levels across different splicing isoforms, even though the site itself does not directly affect splicing motif. Using AlphaFold3, we further explored the differential functional consequences of this site at the protein structural level. In contrast, for the *Vps13* gene, which also contains a single editing site but exhibits minimal sequence differences among isoforms, editing levels of distinct transcripts were highly variable, whereas pooled mature mRNA editing levels remained stable.

For the *stj* gene, which contains clustered editing sites, transcript-specific editing differences are also evident, whereas editing levels across sites in the *qvr* gene show relatively limited isoform-specific variation. Notably, co-editing patterns among multiple sites are generally consistent across transcripts, while a subset of sites exhibits correlations between editing levels and transcript expression level.

Our results demonstrate that RNA editing levels can differ remarkably among isoforms of a single gene. This is particularly evident in genes such as *Adar* which play essential physiological roles, or in genes with neuronal functions which are tightly regulated by RNA editing. These findings suggest that gene-level editing estimates derived from bulk RNA-seq may not accurately capture the functional impact of RNA editing. Different transcript isoforms may serve distinct biological functions and be differentially regulated by RNA editing under varying environmental conditions. From a broader perspective, isoform-specific editing highlights the precision and flexibility of A-to-I RNA editing regulation, contributing to the ability of organisms to adapt to complex and dynamic environments.

In neuronal systems, where rapid and precise modulation of gene function is essential [55], the combination of alternative splicing and RNA editing may provide a multilayered regulatory mechanism. Isoform-specific editing enables fine-tuning of protein function without requiring changes in gene expression levels, offering a rapid and flexible strategy for neural adaptation.

The RT-nPCR and iPrimer PCR approaches complement long-read sequencing technologies by providing highly accurate quantification of RNA editing at isoform resolution, thereby bridging a gap between transcript structure identification and functional modification analysis. Given that dysregulation of RNA editing has been implicated in neurological disorders and cancer [56–58], the ability to resolve isoform-specific editing patterns may uncover previously hidden layers of disease-associated transcriptomic variation.

From a mechanistic perspective, isoform-specific differences in editing levels raise the question of whether RNA editing differentially affects the splicing process of individual transcripts. Splice site prediction analysis indicates that editing sites in *Adar* and *stj*, despite exhibiting strong isoform-specific differences, are not located within canonical splice sites, and nucleotide changes at these positions do not alter predicted splicing outcomes. Previous studies have shown that splicing efficiency can influence RNA editing levels by affecting nuclear retention and RNA processing dynamics, thereby contributing to discrepancies between pre-mRNA and mature mRNA editing levels [48]. Therefore, differences in splicing efficiency may represent one potential mechanism underlying isoform-specific editing levels. In addition, Adar proteins can regulate splicing independently of their catalytic activity [59], and isoform-specific interactions between Adar and pre-mRNA may further contribute to differential splicing outcomes and editing levels.

Despite successfully isolating isoforms followed by quantifying editing levels using Sanger sequencing, our approach remains limited by low throughput. Moreover, because isoform-specific amplification and analysis were performed on a gene-by-gene basis, the global landscape of isoform-specific RNA editing and its regulation by environmental factors remains unclear. Moreover, the experimental parameters of iPrimer PCR, particularly for UTR isoform separation, may require further optimization.

Nevertheless, the RT-nPCR and iPrimer PCR strategies developed in this study provide a relatively low-cost and technically accessible approach for isoform-specific mRNA amplification. Beyond their application in RNA editing analysis, these methods may have broader utility in transcript-level studies, which warrants further exploration. In summary, our works provide new perspectives about the interplay between RNA editing and alternative splicing, and reveal the complex molecular diversity conferred by post-transcriptional mechanisms.

## Materials and Methods

### Fly collection and head RNA extraction

Adult *Drosophila melanogaster* (3-5 days old) were divided into three groups: control, high-temperature, and starvation. Each group included both *w^1118^* and *yw* strains, and male and female flies were processed separately for head RNA extraction.

Prior to RNA extraction, control flies were maintained at 25℃ on standard medium. The medium was prepared by dissolving 8.0 g agar and 20 g sucrose in 400 mL water, followed by sterilization at 120 ℃ for 20 min. The high-temperature group was maintained at 30℃ for 48 h on the same medium. The starvation group was maintained for 48 h on nutrient-free medium prepared with 8.0 g agar in 400 mL water.

Flies were anesthetized using CO_2_ and examined under a stereomicroscope. Male and female flies were separated into different centrifuge tubes (∼50 flies per tube). After allowing recovery from anesthesia, flies were rapidly frozen in liquid nitrogen. Once fully frozen, tubes were removed and sharply tapped to mechanically separate heads from bodies. Heads were then collected using a sieve, immediately transferred into centrifuge tubes, and stored in liquid nitrogen.

Total RNA was extracted from fly heads using the PureLink^TM^ RNA Mini Kit (Invitrogen) according to the manufacturer’s instructions. Extracted RNA was immediately reverse-transcribed into cDNA using the PrimeScript^TM^ RT Reagent Kit with gDNA Eraser (Takara), following the manufacturer’s protocol.

### Known lists of RNA editing sites in *D. melanogaster*

Known lists of RNA editing sites in *D. melanogaster* were collected from multiple previous literatures [33, 40, 47, 60, 61]. We first selected some well-known neuron-related recoding genes that are highly expressed in the *Drosophila* brain. According to the latest *D. melanogaster* genome annotation, genes with only one single transcript isoform were not considered. We prioritize our analysis to the recoding sites with relatively high editing level (between 15% and 90%). The recoding sites must be located in the shared region of different transcript isoforms. Synonymous sites adjacent to the target recoding sites were also considered. Genes *Adar*, *stj*, *qvr*, *Vps13*, and *jef* were selected.

### Visualization of gene model

Gene structure annotations for *Drosophila* genes were obtained from the “Genomic regions, transcripts, and products” section of the NCBI (https://www.ncbi.nlm.nih.gov/) gene database. GFF3 files were downloaded via the “Download Track Data” option and imported into TBtools-II (v2.435) [62]. Gene structure diagrams for individual transcript isoforms were generated using Graphics, BioSequence Structure Illustrator, Gene Structure View. Sequence features surrounding RNA editing sites were examined and visualized using SnapGene 7.1.2 (https://www.snapgene.com/). Schematic illustrations were created using Adobe Illustrator 2024. RNA secondary structures were predicted using ViennaRNA Web Services (http://rna.tbi.univie.ac.at/) [63]. Due to input length limitations, sequences exceeding 10,000 nt were truncated to 10,000 nt segments with the editing site positioned as centrally as possible for structure prediction.

### Primer design and synthesis

Gene and transcript sequences for *Drosophila melanogaster Adar*, *Vps13*, *qvr*, and *stj* were retrieved from the NCBI database as FASTA files. These sequences were imported into SnapGene 7.1.2 for visualization and primer design. Exon-exon junctions suitable for primer placement were identified using the NCBI Genome Data Viewer. Primers for RT-nPCR and RT-qPCR were manually designed for each transcript isoform (**Supplementary Tables S1-S2**).

Primer specificity and melting temperatures (Tm) were validated using NCBI Primer-BLAST, and secondary structures (self-dimers, heterodimers, and hairpins) were evaluated using the Integrated DNA Technologies (IDT) OligoAnalyzer (https://www.idtdna.com/calc/analyzer/). All primers were synthesized by Sangon Biotech using HAP purification.

### Quantification of RNA editing levels across transcript isoforms

Due to the large number of *Adar* isoforms in *Drosophila*, individual isoforms could not be distinguished using a single primer pair. Therefore, reverse transcription nested PCR (RT-nPCR) was employed using cDNA as a template. Based on RNA sequences of 15 *Adar* isoforms obtained from NCBI, multiple primer pairs spanning exon-exon junctions were designed. Sequential rounds of PCR amplification were performed using different primer pairs. Between each round, PCR products were diluted 100-fold and used as templates for subsequent amplification to obtain high concentrations of target isoform products (**Figure 1A**, with primer details in **Supplementary Figure S1**). Amplified products were subjected to Sanger sequencing, and editing levels were quantified based on the proportion of G peaks at the editing sites.

PCR reactions were prepared using EmeraldAmp Max PCR Master Mix (Takara). Products from each round were diluted 100-fold before use as templates for the next round. Final PCR products were sequenced by Tsingke Biotech. Sequencing chromatograms were analyzed using SnapGene 7.1.2, and RNA editing levels were calculated as the proportion of G signal relative to the combined A and G peaks at each editing site (**Supplementary Tables S3-S7**). Data visualization and statistical analyses were performed in R version 4.4.3.

### Prediction of Adar protein structure and RNA binding

Protein sequences of *Drosophila melanogaster Adar* isoforms were downloaded from NCBI in FASTA format and imported into SnapGene 7.1.2 for annotation. Protein domain architecture was analyzed using the NCBI Conserved Domain Search (https://www.ncbi.nlm.nih.gov/Structure/cdd/wrpsb.cgi/) tool and recorded in SnapGene. Amino acid sequence alignments were performed using MEGA version 12 [64] for visualization. Protein tertiary structures were predicted using AlphaFold3(https://alphafoldserver.com/) [65]. The resulting model_0.cif files were visualized in PyMOL 3.1.3.1 (https://pymol.org/), where protein domains, the S>G editing site, and RNA-binding regions were annotated.

### Co-editing analysis of sites in *qvr* and *stj*

Following preprocessing of raw RNA editing data for *qvr* and *stj*, a complete site-by-sample matrix was constructed. Missing values (NA) were processed using a mean imputation strategy, replacing missing entries with the average editing level of the corresponding site across other samples. Pearson correlation analysis based on the completed matrix revealed co-editing relationships among sites. Heatmap visualization highlighted strongly positively correlated site pairs (colored green), while independent site pairs were indicated in white. Clustering analysis based on silhouette coefficients partitioned sites into co-editing modules, within which RNA editing events were highly coordinated. All analyses and visualizations were performed in R version 4.4.3.

### Quantification of transcript isoform expression proportions

RT-qPCR experiments were performed using TB Green Premix Ex Taq^TM^ II (Tli RNaseH Plus, Takara). Ct values for each transcript isoform were measured and used for downstream quantitative analysis (**Supplementary Table S8**). Ct values were converted into relative expression levels using the 2^(−Ct) method. As the objective was to determine the relative proportions of transcript isoforms within each gene, no internal reference gene normalization was applied. Instead, the proportion of each isoform was calculated relative to the total expression of all isoforms within the same sample. Processed data were visualized as stacked bar plots using the ggplot2 package in R to illustrate the relative distribution of transcript isoforms across samples of different sexes. Due to limited sample availability, only control samples were included in RT-qPCR analyses.

### Prediction of whether editing sites directly affect splicing factor recognition

To evaluate whether RNA editing sites directly influence splicing factor recognition through sequence alteration, we employed the Berkeley *Drosophila* Genome Project (BDGP) splice site prediction tool (https://www.fruitfly.org/seq_tools/splice.html/) [66]. Pre-mRNA sequences were first submitted to obtain baseline predictions of splice sites and corresponding scores. Subsequently, the adenosine (A) at each editing site was computationally substituted with guanosine (G), and the modified sequences were reanalyzed. Differences in predicted splice sites and scores before and after substitution were compared to assess whether RNA editing could directly modulate splicing through sequence changes.

### Quantification of editing levels in UTR isoforms using interfering-Primer PCR

Interfering-Primer PCR (iPrimer PCR) is a strategy designed to enable selective amplification of a target transcript in the presence of closely related longer transcripts that differ only by an extended terminal sequence. This is achieved through the introduction of an interfering primer (iPrimer) that selectively suppresses amplification of the longer, non-target transcript (**Figure 1B**).

First, primers were designed to anneal to the terminal region of the target transcript and to an exon-exon junction on the opposite side of the editing site, enabling amplification of the target cDNA. The iPrimer was then designed with the following considerations: to prevent binding of the terminal amplification primer to the longer non-target transcript while avoiding binding to the target transcript itself, the 3’ end (∼10 nt) of the iPrimer was designed to match the sequence near the terminal region of the target-specific primer-binding site. The 3’ end of the iPrimer was phosphorylated to prevent extension.

To ensure preferential binding to the longer non-target transcript prior to amplification primer annealing, the 5’ region of the iPrimer included a sequence present in the longer transcript but absent in the shorter target transcript. This design increased the overall length and melting temperature (Tm) of the iPrimer, typically at least 10℃ higher than that of the amplification primers, allowing it to bind at higher temperatures.

During PCR cycling, an Inhibition phase was introduced immediately after denaturation, allowing sufficient time for iPrimer binding to non-target transcripts. This was followed by a Stabilization phase, set at a temperature approximately 3℃ below the iPrimer Tm (but still > 10℃ above the amplification primer Tm), to maintain stable iPrimer binding prior to conventional annealing and extension steps (**Supplementary Figure S7**). Following iPrimer PCR, products were subjected to Sanger sequencing. An additional internal sequencing primer (not used for amplification) was designed to ensure that the editing site fell within the effective read length.

Design principles for iPrimers include:

1. Tm at least 10℃ higher than amplification primers.
2. 3’ end phosphorylation to prevent extension.
3. ∼10 nt at the 3’ end corresponding to the truncated terminal sequence of the short transcript.
4. 5’ sequence derived from the extended region unique to the longer transcript.
5. iPrimer concentration higher than that of individual amplification primers.

By combining iPrimer PCR with quantitative PCR (iPrimer qPCR), relative expression levels of “embedded” transcript isoforms can be quantified. Due to potential primer-primer interactions, especially in the presence of long iPrimers, the temperatures of the Inhibition and Stabilization phases were slightly increased compared to standard PCR conditions. Reaction components for short transcript amplification were maintained identical to those for long transcripts, except for the inclusion of iPrimer.

Because qPCR amplicon length must be limited, amplification of NM_001259394.2 inevitably included some pre-mRNA, resulting in slight overestimation. However, independent quantification of pre-mRNA versus mature mRNA indicated that this effect was minimal and did not alter the observed dominance of the long UTR isoform NM_001259394.2 over NM_137139.2.

iPrimer PCR reactions were prepared using EmeraldAmp Max PCR Master Mix (Takara), and iPrimer qPCR reactions were conducted using TB Green Premix Ex Taq^TM^ II (Tli RNaseH Plus, Takara) (**Supplementary Table S9**). Primers were designed using SnapGene 7.1.2 (**Supplementary Tables S1-S2**). PCR products were sequenced by Tsingke Biotech, and RNA editing levels and expression data are provided in **Supplementary Tables S7-S8**.

Motif and regulatory element prediction for UTR isoforms was performed using RegRNA 3.0 [67].

### Quantification of editing levels in UTR isoforms using circularized cDNA

To validate the results obtained by iPrimer PCR, we additionally employed a circularized cDNA approach to isolate and sequence short transcript isoforms (**Figure 1C**). Purified cDNA from *Drosophila* head samples was first diluted to 1-5 ng/μL. At low DNA concentrations, intermolecular collisions are reduced, while intramolecular ligation is favored, thereby increasing circularization efficiency [68].

The diluted cDNA was subjected to self-ligation using T4 DNA ligase. To selectively enhance circularization efficiency of the target short transcript, a splint oligonucleotide complementary to both ends of the target cDNA was introduced. This splint bridges the two ends to form a transient double-stranded structure, facilitating efficient ligation by T4 ligase [69].

Following circularization, target short transcripts and longer non-target transcripts generate distinct sequences at the circularization junction. Linear cDNA was subsequently removed using T5 exonuclease. Primers specific to the circularization junction of the short transcript were then designed for inverse PCR, enabling selective amplification of the target isoform. Nested PCR was further employed to increase product yield for sequencing.

To further validate editing levels in short transcript isoforms, we employed a circularized cDNA strategy to generate unique junction sites for specific amplification.

First, cDNA extracted from fly heads was purified using the EasyPure PCR Purification Kit (TransGen Biotech). Briefly, 5 μL of cDNA was mixed with 25 μL Binding Buffer, loaded onto a spin column, incubated for 1 min, and centrifuged at 14,000 rpm for 1 min. The flow-through was discarded. The column was washed with 65 μL Wash Buffer, incubated for 1 min, and centrifuged at 14,000 rpm for 1 min. After discarding the flow-through, the column was centrifuged again at 14,000 rpm for 2 min to remove residual buffer. During this step, 35 μL Elution Buffer was preheated to 60℃. The column was then transferred to a clean tube, and cDNA was eluted with 5 μL Elution Buffer after 1 min incubation at room temperature. The purified cDNA was quantified and diluted to 1-5 ng/μL.

Circularization was performed using T4 DNA Ligase (New England Biolabs), with reaction components prepared according to the manufacturer’s instructions (**Supplementary Tables S9**). The reaction mixture was first incubated at room temperature for 5 min to facilitate splint oligonucleotide bridging. Subsequently, ligation was carried out at 16℃ for 16 h. During this period, the water bath was turned off for 10 min every 1.5 h to prevent overheating and to enhance splint-cDNA hybridization. After ligation, the enzyme was inactivated at 65℃ for 10 min.

Linear cDNA was removed using T5 Exonuclease (New England Biolabs), following the manufacturer’s instructions (**Supplementary Tables S9**). The reaction was incubated at 37℃ for 30 min to degrade linear cDNA, followed by heat inactivation at 70℃ for 10 min and cooling on ice for 2 min.

To enrich circularized products, the reaction mixture was purified again using the EasyPure PCR Purification Kit. Briefly, 50 μL of sample was mixed with 250 μL Binding Buffer, applied to a spin column, incubated for 1 min, and centrifuged at 14,000 rpm for 1 min. After discarding the flow-through, the column was washed with 650 μL Wash Buffer and centrifuged under the same conditions. The column was then centrifuged empty for 2 min to remove residual buffer. Meanwhile, 320 μL Elution Buffer was preheated to 60℃. cDNA was eluted with 50 μL Elution Buffer after 1 min incubation at room temperature.

Inverse PCR was then performed to specifically amplify the circularized short transcript isoforms. Reactions were prepared using EmeraldAmp Max PCR Master Mix (Takara), with a higher template input than standard PCR to enhance product yield (**Supplementary Tables S9**). The inverse PCR products were diluted 100-fold and subjected to nested PCR under the same reaction conditions as described above.

Final PCR products were sequenced by Tsingke Biotech using Sanger sequencing to determine RNA editing levels. Primers used for inverse PCR and nested PCR were designed using SnapGene 7.1.2 (see **Supplementary Tables S1**). Editing level data are provided in **Supplementary Tables S7**.

## Supporting information

Contains Supplementary Figures 1-7 and the titles of Supplementary Tables 1-9.

Contains Supplementary Tables 1-9 in excel format.

## Abbreviations

A-to-I: adenosine-to-inosine.
CDS: coding sequence.
iPrimer PCR: interfering-Primer
PCR. NGS: next-generation sequencing.
RT-nPCR: reverse transcription nested
PCR. UTR: untranslated region.

## Acknowledgements

We thank the High-performance Computing Platform of China Agricultural University platform for the computational support.

## Data availability

All raw Sanger sequencing files used in this study have been deposited to Figshare under link: https://doi.org/10.6084/m9.figshare.32049537. Relevant processed data have been integrated into the **Supplementary Tables** of this manuscript.

## Authors’ contributions

Conceptualization & supervision: Y.D. and H.L.

Writing – original draft: Y.D., Z.Y., W.C., and H.L.

Writing – review & editing: Y.D., Z.Y., W.C., and H.L.

All authors approved the submission of this manuscript.

## Declarations

### Ethics approval and consent to participate

Not applicable.

### Consent for publication

Not applicable.

