## Supplementary material for "RT-nested and interfering-Primer PCR reveal prevalent isoform-specific A-to-I RNA editing in neuronal genes": Contains Supplementary Figures 1-7 and the titles of Supplementary Tables 1-9.

### Supplemental Figures

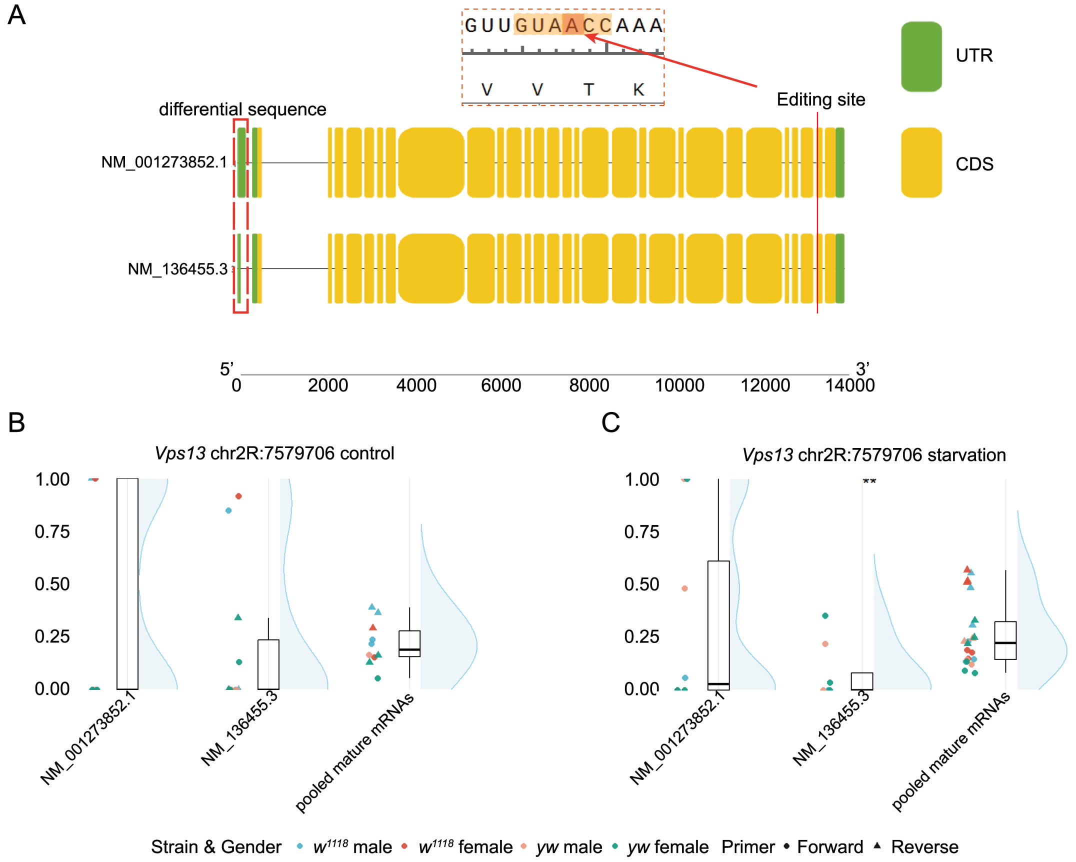


**Supplementary Figure S1. Gene structure and RNA editing levels of the *Vps13* gene.** (A) Gene models and editing site positions of the two *Vps13* transcripts. The sequences of the two transcripts are highly similar. Green indicates UTR regions, and yellow indicates CDS regions. Transcript structures were visualized by importing GFF3 files downloaded from NCBI into TBtools-II (v2.435). (B-C) Editing levels at site chr2R:7579706 in the two *Vps13* transcripts and pooled mature mRNA. Pre-mRNA editing levels are not available. (B) is for control group and (C) is for starvation group. The high-temperature group was not analyzed due to insufficient sample amount. Color scheme was identical to that of the *Adar* gene.

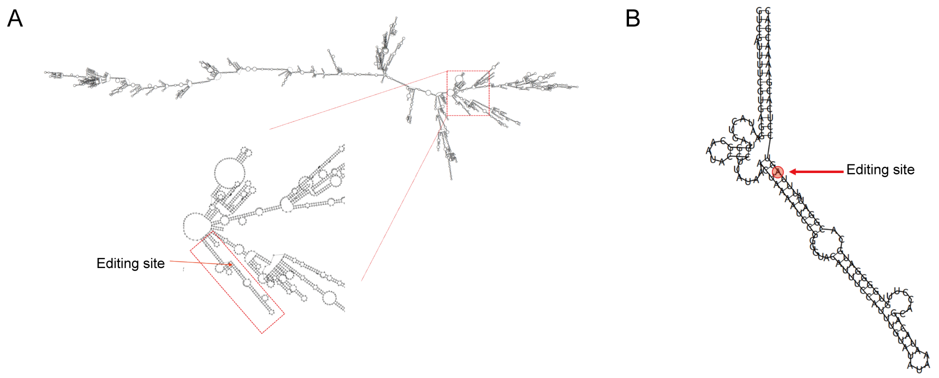


**Supplementary Figure S2. A conserved region from *Adar* pre-mRNA (around S>G recoding site) used for AlphaFold analysis on protein-RNA interaction.** (A) Full-length pre-mRNA structure predicted using ViennaRNA Web Services. (B) Folding the local sequence produces a nearly identical secondary structure.

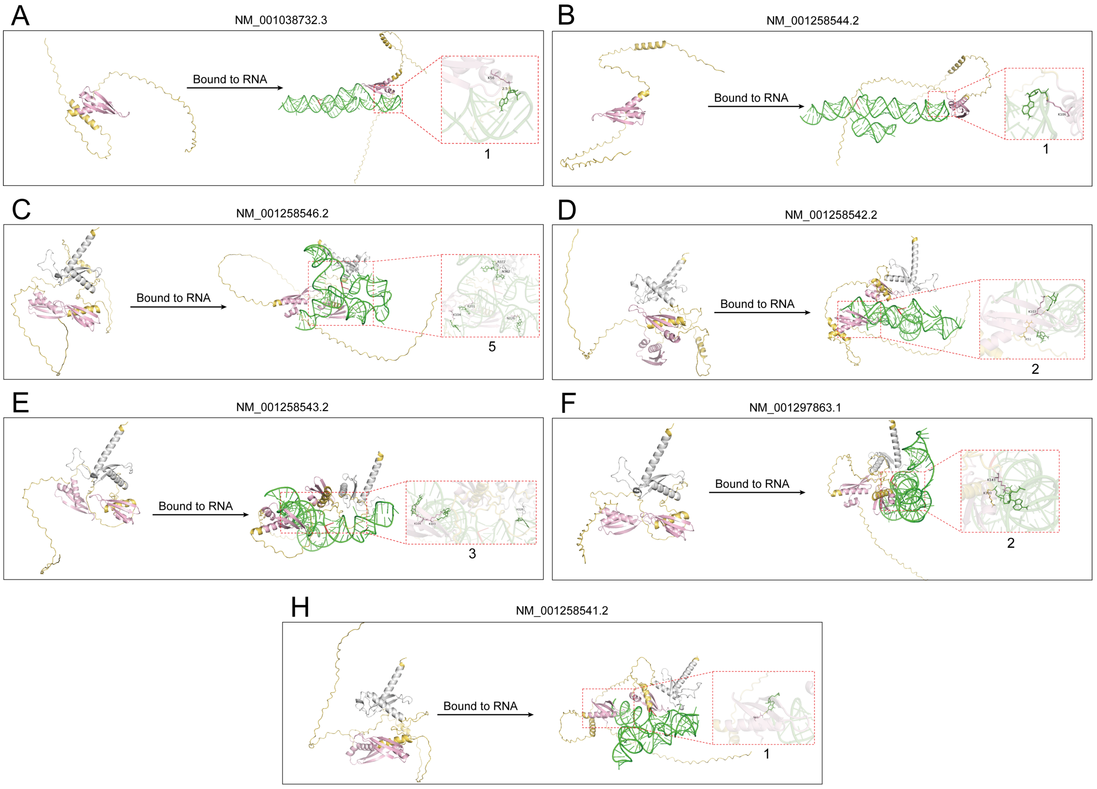


**Supplementary Figure S3. AlphaFold3-predicted interactions between RNA and proteins encoded by transcripts lacking or partially lacking the catalytic domain.** (A-B) Adar proteins lacking the ADEAMc domain can bind RNA via the remaining DSRM_STRBP_RED-like_rpt1 domain but are unable to unwind dsRNA structures. (C-H) Adar proteins containing incomplete ADEAMc domains retain the ability to bind RNA and unwind dsRNA, indicating partial functional capacity. Visualization was performed using PyMOL 3.1.3.1。

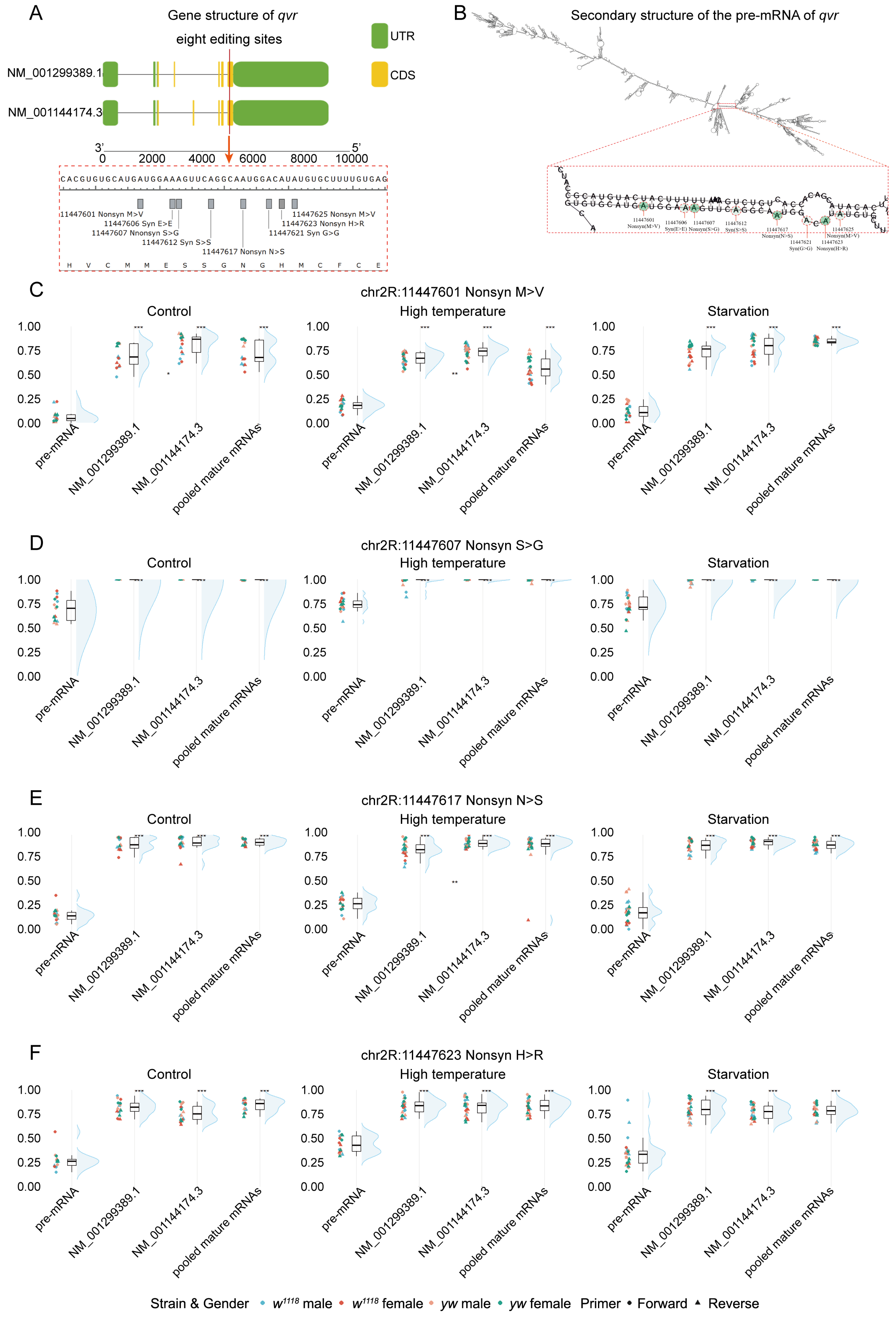


**Supplementary Figure S4. Clustered editing sites in the *qvr* gene.** (A-B) Transcript structures and editing site coordinates of *qvr*. (C-F) RNA editing levels measured by Sanger sequencing. Color scheme was identical to that of the *Adar* gene.

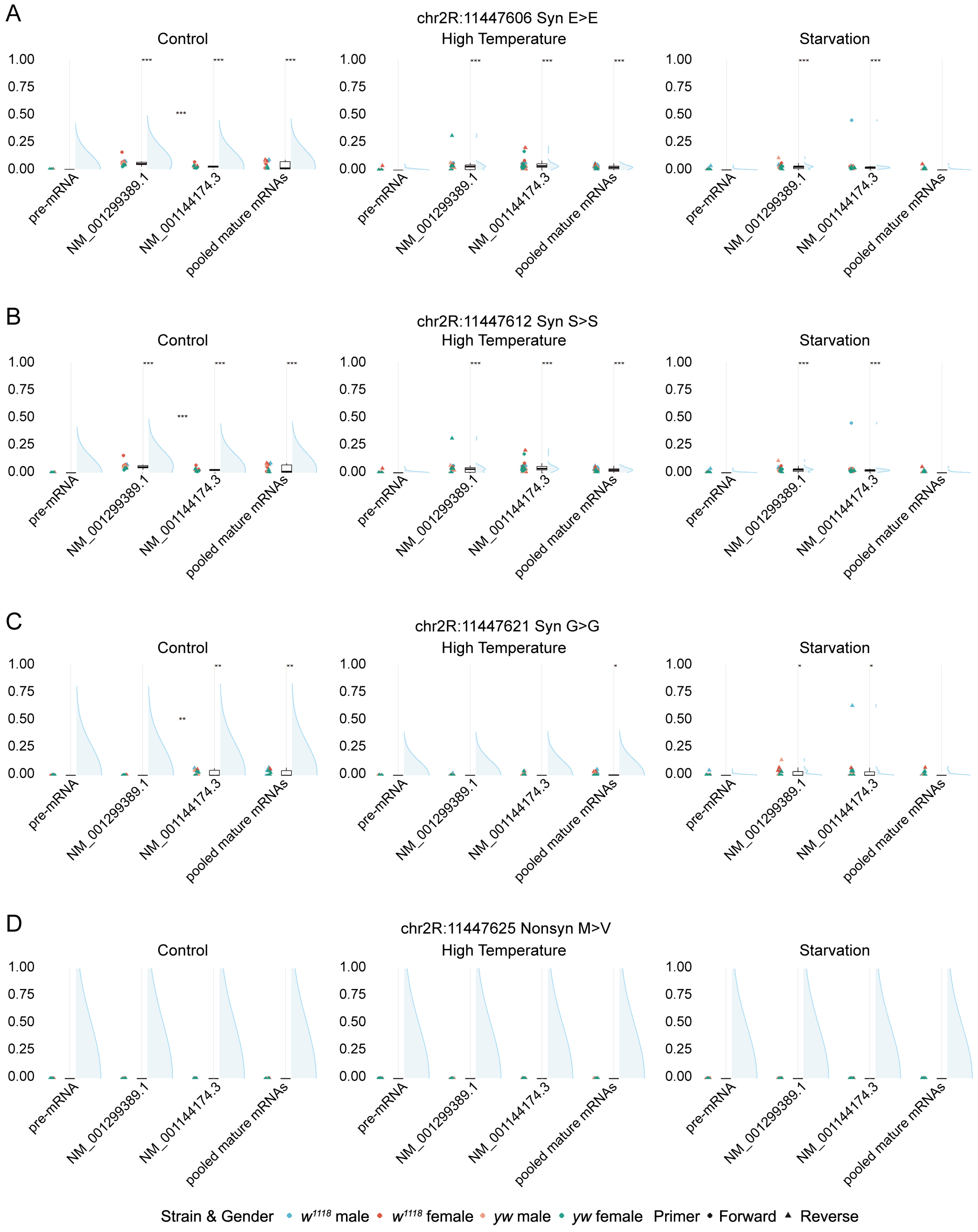


**Supplementary Figure S5. Demonstration of editing levels approaching zero at specific *qvr* sites.** (A-D) Editing levels at four sites (chr2R:11447606, 11447612, 11447621, and 11447625) are close to zero across all transcripts.

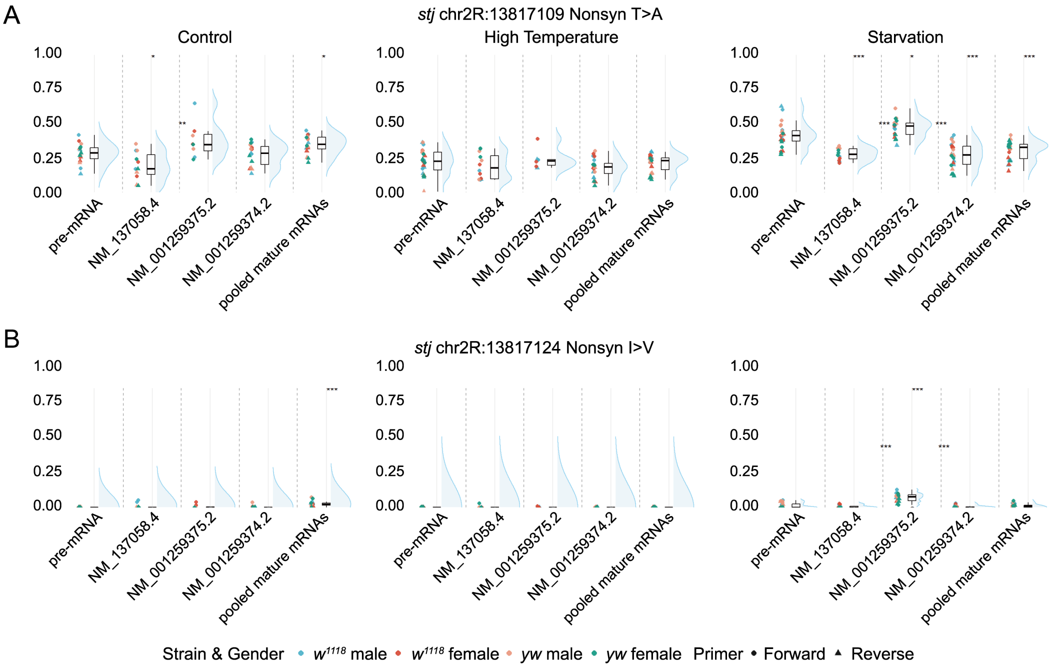


**Supplementary Figure S6. Editing levels at two *stj* gene sites.** (A) Editing levels at chr2R:1381710 vary among transcripts. (B) Editing level at chr2R:13817124 is zero.

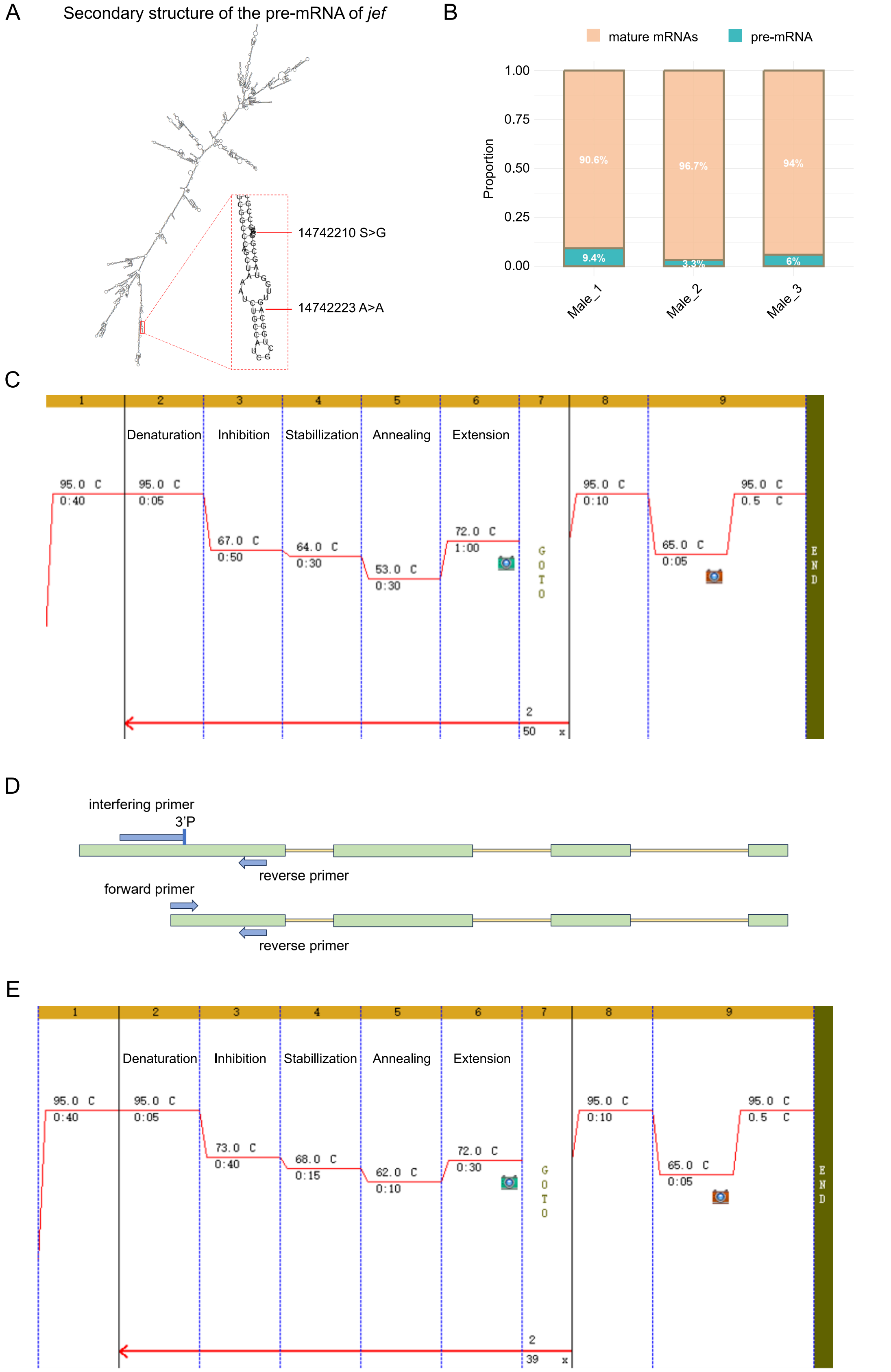


**Supplementary Figure S7. Isoform-specific amplification of the *jef* gene using iPrimer PCR.** (A) Distribution of editing sites within *jef* pre-mRNA. RNA structure was predicted using ViennaRNA Web Services. (B) RT-qPCR analysis of pre-mRNA and mature mRNA. Due to limited sample amount, only control $w^{1118}$ males were analyzed. (C) iPrimer PCR cycling conditions. An additional inhibition step was introduced after each denaturation step to facilitate binding of iPrimer to non-target long transcripts. A subsequent stabilization step at lower temperature enhances binding stability before standard annealing and extension steps. (D) Schematic of iPrimer qPCR. iPrimer prevents amplification of long transcripts, enabling specific amplification of target isoforms. Amplicon length was designed to meet qPCR requirements. (E) iPrimer qPCR cycling conditions. Temperatures for inhibition and stabilization steps were slightly increased to reduce primer dimer formation.

### Supplementary Tables

**Supplementary Table S1.** Primers used for isoform-specific amplification by RT-nPCR or iPrimer PCR and subsequent Sanger sequencing of RNA editing levels.

**Supplementary Table S2.** Primers used for RT-qPCR quantification of transcript expression ratios.

**Supplementary Table S3.** RNA editing levels of the *Adar* gene across transcripts, treatments, strains, and sexes.

**Supplementary Table S4.** RNA editing levels of the *Vps13* gene across transcripts, treatments, strains, and sexes.

**Supplementary Table S5.** RNA editing levels of the *qvr* gene across transcripts, treatments, strains, and sexes.

**Supplementary Table S6.** RNA editing levels of the *stj* gene across transcripts, treatments, strains, and sexes.

**Supplementary Table S7.** RNA editing levels of the *jef* gene across transcripts, treatments, strains, and sexes.

**Supplementary Table S8.** Raw Ct values from RT-qPCR measurements of transcript expression ratios.

**Supplementary Table S9.** Composition and proportions of reaction mixtures used for specific cDNA amplification.
